# DECODER: A probabilistic approach to integrate big data reveals mitochondrial Complex I as a potential therapeutic target for Alzheimer’s disease

**DOI:** 10.1101/302737

**Authors:** Safiye Celik, Josh C. Russell, Cezar R. Pestana, Ting-I Lee, Shubhabrata Mukherjee, Paul K. Crane, C. Dirk Keene, Jennifer F. Bobb, Matt Kaeberlein, Su-In Lee

## Abstract

Identifying gene expression markers for Alzheimer’s disease (AD) neuropathology through meta-analysis is a complex undertaking because available data are often from different studies and/or brain regions involving study-specific confounders and/or region-specific biological processes. Here we introduce a novel probabilistic model-based framework, DECODER, leveraging these discrepancies to identify robust biomarkers for complex phenotypes. Our experiments present: (1) DECODER’s potential as a general meta-analysis framework widely applicable to various diseases (e.g., AD and cancer) and phenotypes (e.g., Amyloid-β (Aβ) pathology, tau pathology, and survival), (2) our results from a meta-analysis using 1,746 human brain tissue samples from nine brain regions in three studies — the largest expression meta-analysis for AD, to our knowledge —, and (3) *in vivo* validation of identified modifiers of Aβ toxicity in a transgenic *Caenorhabditis elegans* model expressing AD-associated Aβ, which pinpoints mitochondrial Complex I as a critical mediator of proteostasis and a promising pharmacological avenue toward treating AD.

## Introduction

Alzheimer’s disease (AD) is the 6^th^ most common cause of death in the United States; however, no effective therapy exists to delay or prevent its onset or progression.^1^ Neuritic plaques and neurofibrillary tangles are two neuropathological hallmarks of AD, whose basic building blocks are Amyloid-β (Aβ) peptide and the tau protein, respectively. Aβ is a 40 to 42 amino acid-peptide, which is generated by proteolytic cleavages of the amyloid precursor protein *(APP)* located on chromosome 21.^2^ While the precise function of *APP* is not known, mutations in this gene are thought to lead to familial susceptibility to AD.^3^ The amyloid cascade hypothesis^4^ posits that aggregation and extracellular deposition of the misfolded Aβ peptide (specifically, the Aβ_1–42_ isoform) plays a causal role in several AD-related events, including neurofibrillary tangle formation, resulting in neuronal death and neurodegeneration.^5^ However, causality remains unproven.^6^ This hypothesis has given rise to several Aβ-centric treatment approaches for AD, such as blocking *APP* cleavage using β-secretase inhibitors, which has been attempted to reduce Aβ formation and deposition, but several concerns have been raised about its efficacy.^7^ Tau, the other hallmark of AD, is a brain-specific, axon-enriched, microtubule-associated protein^8^ encoded by a gene located on chromosome 17. It is phosphorylated post-translationally and has phosphorylation levels that are significantly higher in brains of AD patients, suggesting that hyper-phosphorylation may lead to tau’s pathogenic role in AD.^9^

At present, we lack well-founded knowledge of the set of genes whose expression affects the formation of neuritic plaques and neurofibrillary tangles and the protective and pathological responses to these purportedly toxic lesions: until recently, we have had limited access to gene expression data and neuropathological phenotypes from post-mortem brain tissues. Biologists have only now begun gathering both gene expression data and Aβ and tau measures from human brain tissues,^10^ providing a new paradigm of system-level AD data for information mining. As part of the Accelerating Medicines Partnership (AMP)-AD project, Sage Bionetworks recently began sharing and integrating multi-dimensional human ‘omic’ data from more than 2,000 human brains autopsied in several studies through the AMP-AD Knowledge Portal.^11^ Of those studies, the Religious Orders Study^12^ and the Memory and Aging Project^13^ (ROSMAP) are longitudinal, clinical-pathologic cohort studies of aging and AD; they provide detailed neuropathology quantifications and omics data, including RNA-Seq, miRNA, DNA methylation, and histone modification measured in the dorsolateral prefrontal cortex of more than 700 brains. The Mount Sinai Brain Bank (MSBB) study provides microarray data and partial neuropathology information from 19 regions of more than 60 brains^14^ and RNA-Seq data from four regions of more than 200 brains. The Adult Changes in Thought (ACT) study —a longitudinal, population-based prospective cohort study of brain aging and dementia— provides detailed neuropathology quantifications and RNA-Seq data from four regions of around 100 autopsied brains.^15^ The increasing number of studies that attempt to obtain brain gene expression and neuropathology data makes meta-analysis (i.e., a statistical analysis combining data from multiple studies to develop a single conclusion) an increasingly powerful approach.

Identifying expression markers for neuropathological phenotypes through the meta-analysis of these studies involves three major challenges. First, expression datasets obtained from different AD studies often present data from different brain regions that contain different cell type compositions, have different functions, and differ in their relevance to AD. While we have a general understanding of how AD progresses across the brain,^16^ it is not of an extent that lets us assign a specific weight value to each brain region in a meta-analysis setting to identify AD neuropathology mechanisms. Second, datasets from different studies involve confounding factors due to differences in subject compositions or experimental procedures used to generate data (i.e., severe batch effects). Finally, the brain tissues used for expression and phenotype profiling do not always match, and each study uses slightly different methods to measure Aβ or tau levels. These factors lead to different sources and levels of noise across studies. Given these challenges, natural questions that arise include whether we can leverage discrepancies across studies or regions to filter out false positive expression markers by focusing on concordant expression associations across different sources.

We present DECODER (<u>d</u>iscov<u>e</u>ring <u>co</u>ncor<u>d</u>ant <u>e</u>xpression ma<u>r</u>kers), a probabilistic model-based framework to identify robust expression markers for a phenotype (e.g., Aβ or tau levels in brain). DECODER adopts a probabilistic model we designed to capture the concordance of gene expression-neuropathology associations across brain regions and/or studies. Interestingly, we show that DECODER’S learning algorithm to estimate the model parameters needs only the expression-phenotype associations (i.e., summary statistics) computed for each dataset rather than the original expression and phenotype data. Specifically, DECODER uses the Pearson’s correlation coefficient (PCC) between an expression level and a phenotype as a *kernel* that projects the original expression and phenotype data into a new gene-phenotype space in which different data sources (e.g., different studies, different tissues, different brain regions, etc.) are more comparable. This is an important advantage that can facilitate meta-analysis, especially when we have access to only summary statistics of expression-phenotype associations, not the original data. We show that concordant expression markers substantially improve statistical and biological consistency compared to expression markers identified from individual studies, tissues, or brain regions considered separately.

Our experimental results can be divided into three main categories. First, using DECODER, we identify genes that are consistently associated with quantified levels of Aβ across multiple human brain regions. This demonstrates the possibility of performing meta-analysis in the highly challenging setting, where datasets involve study-specific confounders or brain region-specific biological processes. Second, by applying DECODER to expression-tau associations in AD and expression-patient survival associations in cancer, we show that DECODER is a powerful general meta-analysis framework widely applicable to various phenotypes or diseases. Third, we experimentally validate the identified expression markers for Aβ in an animal model of Aβ proteotoxicity *in vivo*. We show that knockdowns of a mitochondrial Complex I gene very highly ranked by DECODER and 12 additional Complex I genes attenuate disease in a nematode genetically modified to express human Aβ_1–42_, which causes an age-related paralysis phenotype. Our experimental results advance scientific knowledge of *C. elegans* biology by revealing functions of previously uncharacterized *C. elegans* genes and a potential mechanism for how Aβ toxicity is regulated in this nematode. More importantly, our results reveal Complex I to be a promising therapeutic target for AD and offer hope for the treatment of this disease.

The development of DECODER and *in vivo* confirmatory experimentation of DECODER’s findings in *C. elegans* address fundamental limitations of the standard computational approach to biomarker discovery, which focuses on genes whose expression levels are statistically associated with a phenotype of interest^3–6^ (e.g., neuropathological phenotypes of neuritic plaques or neurofibrillary tangles). Unfortunately, false positive findings are very common in this approach, as evidenced by the low success rates (less than 1%) of replication in independent datasets and translation to clinical practice.^3–6^ The high false positive rate indicates two challenges posed by the current approach. First, high dimensionality, hidden variables, and feature correlations create a discrepancy between statistical associations and true biological interactions; DECODER provides *a new feature selection criterion* to filter out false positive associations by focusing on concordant expression-phenotype associations. Second, correlational results from observational data without *supporting results from interventional experiments* do not prove causal associations. This paper presents an integrative framework that aims to resolve both challenges.

In recent years, significant research has focused on identifying genes that influence the risk of developing, age of onset, and progression of AD. Mutations in genes *APP, PSEN1*, and *PSEN2* are known to cause familial early-onset AD.^17^ In addition, several bona fide genes for late-onset sporadic AD have been identified by genome-wide association studies (GWAS), including *APOE, CLU, ABCA7, SORL1, PICALM, PLD3*, and *ADAM10*.^18^ However, those genetic modifiers account for only a small fraction of AD risk^19,20^ and provide no clear picture of AD neuropathology mechanisms. With the growing availability of expression and neuropathological phenotype data, DECODER will facilitate the pace of discovery in molecular mechanisms of AD pathogenesis.

## Results

This work seeks to identify robust expression markers for disease phenotypes, specifically expression markers for the Aβ neuropathological phenotype, which could be used as therapeutic targets for individuals with AD. We leverage concordance of expression associations with neuropathological phenotypes to filter out noise and extract information likely to be enriched for true signals from *big* high-throughput brain tissue data. We hypothesized that while varied brain regions are affected by AD throughout the progression of the disease and each of these regions is affected with different severities across individuals in the same stage of AD, the basic molecular mechanisms driving the development of and response to neuropathology might be common across regions. Thus, we believe that genes whose associations with neuropathology are concordant in multiple brain regions and studies are more likely to be true molecular markers, and we can use association concordance to reduce the dimensionality of gene expression data into a smaller set of highly informative genes.

Our experiments use data from the ACT study and the AMP-AD RNA-Seq studies, which quantify neuropathology findings.^11^ Thus, we draw data from a total of nine brain regions and 1,746 post-mortem brain tissue samples described in three different studies — ROSMAP,^12,13^ ACT,^15,21^ and MSBB. Our analysis included 14,912 protein-coding genes that have a nonzero RNA-Seq read count in at least one-third of the samples in each study and overlapping across the three studies. The nine regions we used and the number of samples that have both gene expression levels and Aβ quantification are listed in Table 1 by brain region. The Methods section provides details about our data collection.

**Table 1.** The nine brain regions from three studies we used in this work, and the corresponding number of samples that have both gene expression and Aβ quantifications and hence were used to compute gene expression-Aβ associations.

| Region acronym<br>(Study-brain region) | Brain region | Sample size |
| --- | --- | --- |
| ROSMAP-DPFCx | Dorsolateral prefrontal cortex | 542 |
| ACT-TCx | Temporal cortex | 85 |
| ACT-PCx | Parietal cortex | 79 |
| ACT-HIP | Hippocampus | 80 |
| ACT-FWM | Forebrain white matter | 81 |
| MSBB-BM10 | Brodmann area 10 (frontopolar cortex) | 244 |
| MSBB-BM36 | Brodmann area 36 (ectorhinal cortex) | 200 |
| MSBB-BM44 | Brodmann area 44 (opercular part of the inferior frontal gyrus) | 208 |
| MSBB-BM22 | Brodmann area 22 (posterior part of the first temporal gyrus) | 227 |

Figure 1 summarizes the DECODER methodology. Based on a probabilistic model we propose (Methods), we introduce three versions of a method to score genes based on their concordant associations with neuropathology levels in multiple brain regions. We refer to genes that were highly ranked by those three concordance-based approaches as “concordant expression markers (CEMs)”. Using multiple statistical and biological evaluation metrics, we investigate the performance of each new scoring system. We demonstrate that all three concordance-based scores are statistically more robust than individual region-based scores. Further, the CEMs are biologically more relevant to AD than the top genes identified based on individual region-based scores. Based on our computational evaluations, we obtained a small subset of genes that are potential drivers for Aβ pathology, and we conducted biological experiments to validate our findings.

**Figure 1.**
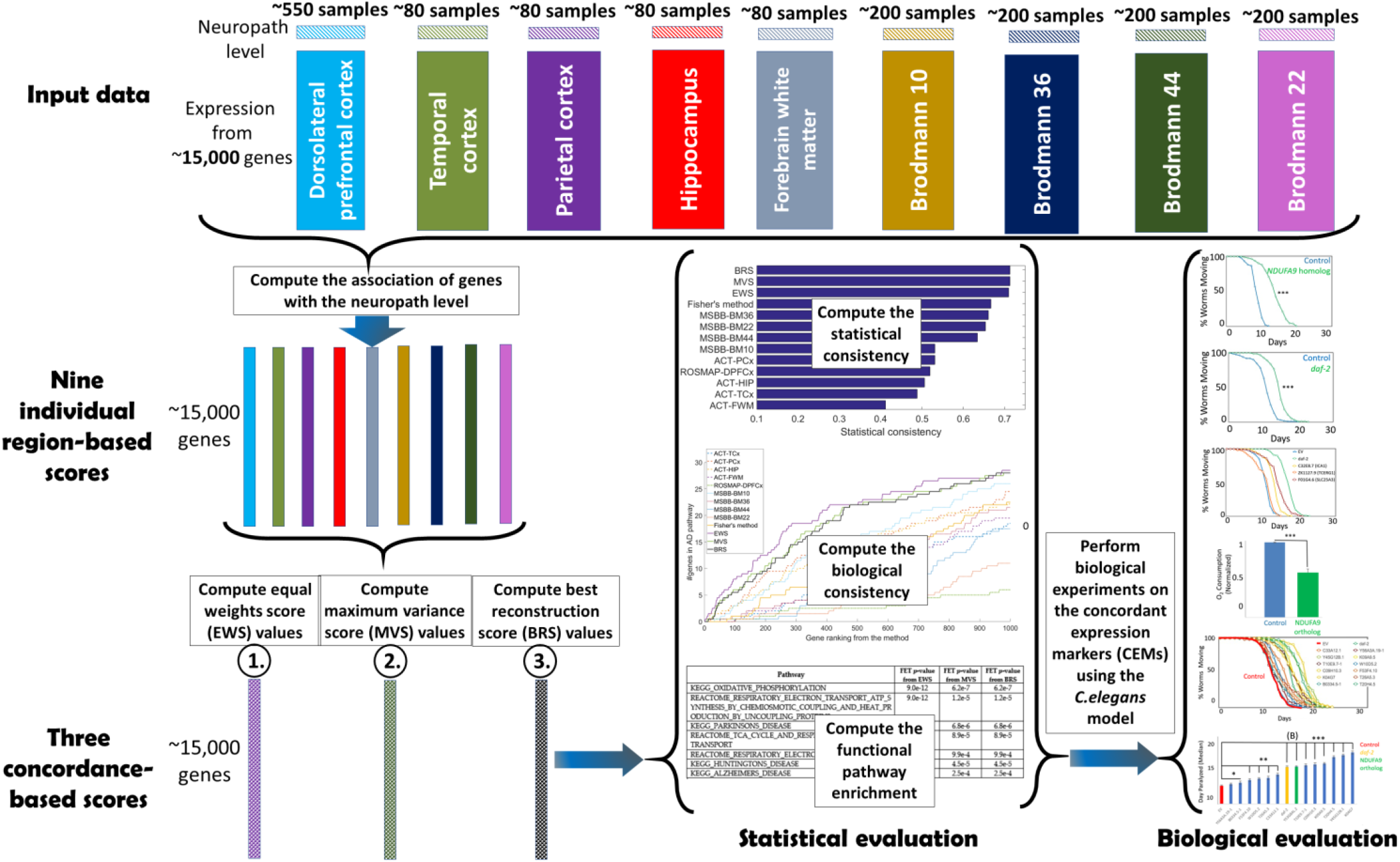
DECODER methodology. Based on a probabilistic model of the observed gene expression and phenotype data, we introduce three different approaches to score the genes based on their concordant associations with neuropathology levels in different brain regions and evaluate them using our statistical and biological evaluation experiments.

Our results, highlighted here, are described in more depth in the subsections that follow:

**Result 1:** Genes associated with neuropathology tend to be concordant across brain regions and studies.
**Result 2:** Brain regions relevant to AD are highly weighted by DECODER.
**Result 3:** Concordance-based approaches identify statistically robust markers of AD pathology.
**Result 4:** CEMs are highly enriched for genes known to be relevant to AD.
**Result 5:** CEMs are highly enriched for pathways relevant to AD.
**Result 6:** Concordance-based approaches are robust to different phenotypes/diseases, including AD tau phenotype and cancer survival.
**Result 7:** The *NDUFA9* gene is biologically validated *in vivo* to be a modifier of Aβ toxicity.
**Result 8:** Knockdown of *NDUFA9*’s nematode homolog strongly decreases whole animal oxygen consumption.
**Result 9:** *In vivo* validation of 12 additional Complex I genes identifies mitochondrial Complex I as a potential AD drug target.
**Result 10:** A multi-omic module subnetwork involving Complex I highlights mechanisms relevant to AD.

### Result 1. Genes associated with neuropathology tend to be concordant across brain regions and studies

We first computed the association of each gene’s expression with Aβ levels in each study. Interestingly, for most pairs of the nine regions, we observed a significant overlap in the top genes whose expression is significantly associated with Aβ levels in that region (Table 2). ACT-FWM was relatively less concordant with the other eight regions, which is not surprising since the other regions contain mostly gray matter expression and neuropathology, and gray and white matter are known to be biologically and functionally distinct.

**Table 2.**
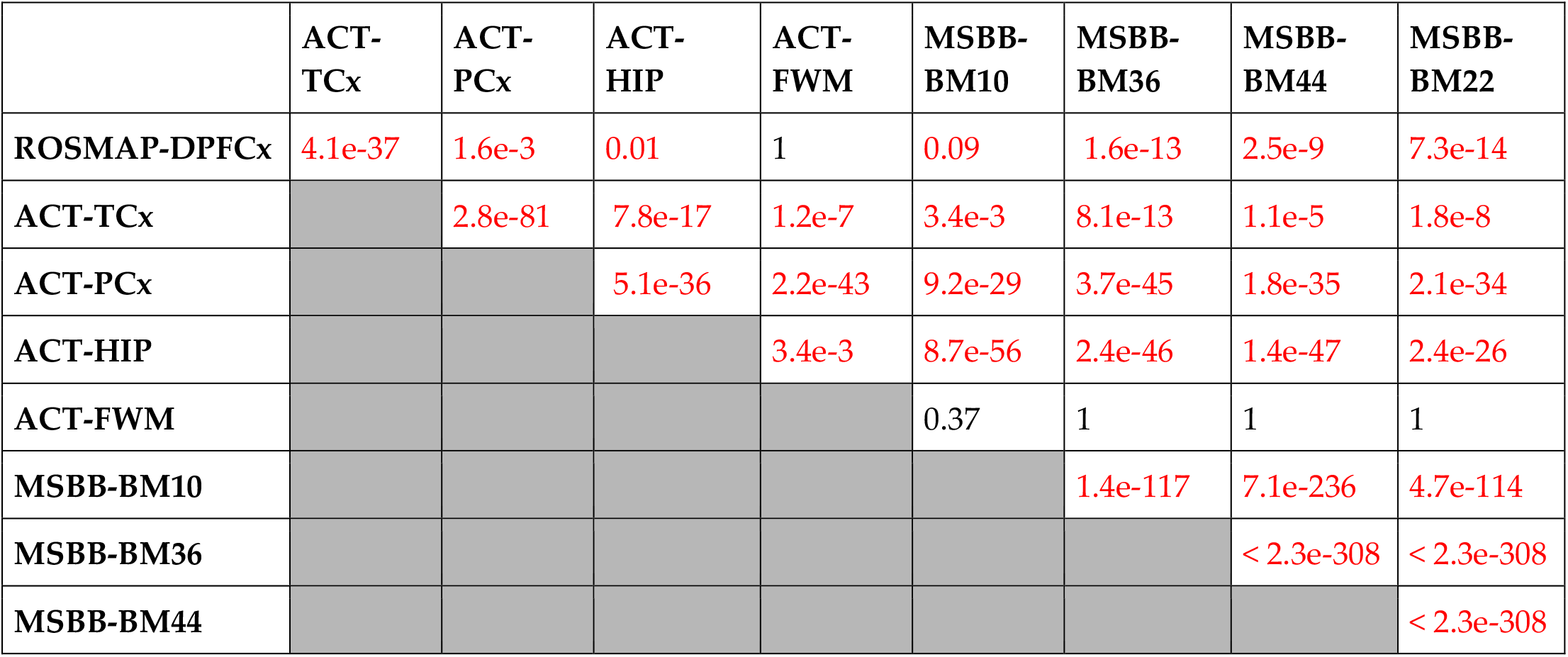
Fisher’s exact test p-value associated with the overlap between the top 1,000 Aβ-associated genes in each region pair (1^st^ column). We highlight in red all pairs that have a statistically significant overlap at p < 0.1.

Figure 2 shows a heatmap of Aβ associations for the ~15,000 genes we included in our analysis. The genes and regions are hierarchically clustered based on their similarities in terms of the strength of association of gene expression with the Aβ phenotype. The rows and columns are rearranged to group similar regions or genes. Bright red and green portions of the heatmaps correspond to the clusters. Although regions from the same study tend to group together, the heatmap shows that MSBB-BM10 (frontopolar cortex) connects to ROSMAP-DPFCx (dorsolateral prefrontal cortex) in the dendrogram before it connects to other regions from the same study. This observation is consistent with the fact that Brodmann area 10 is in close proximity to the dorsolateral prefrontal cortex in the brain.

**Figure 2.**
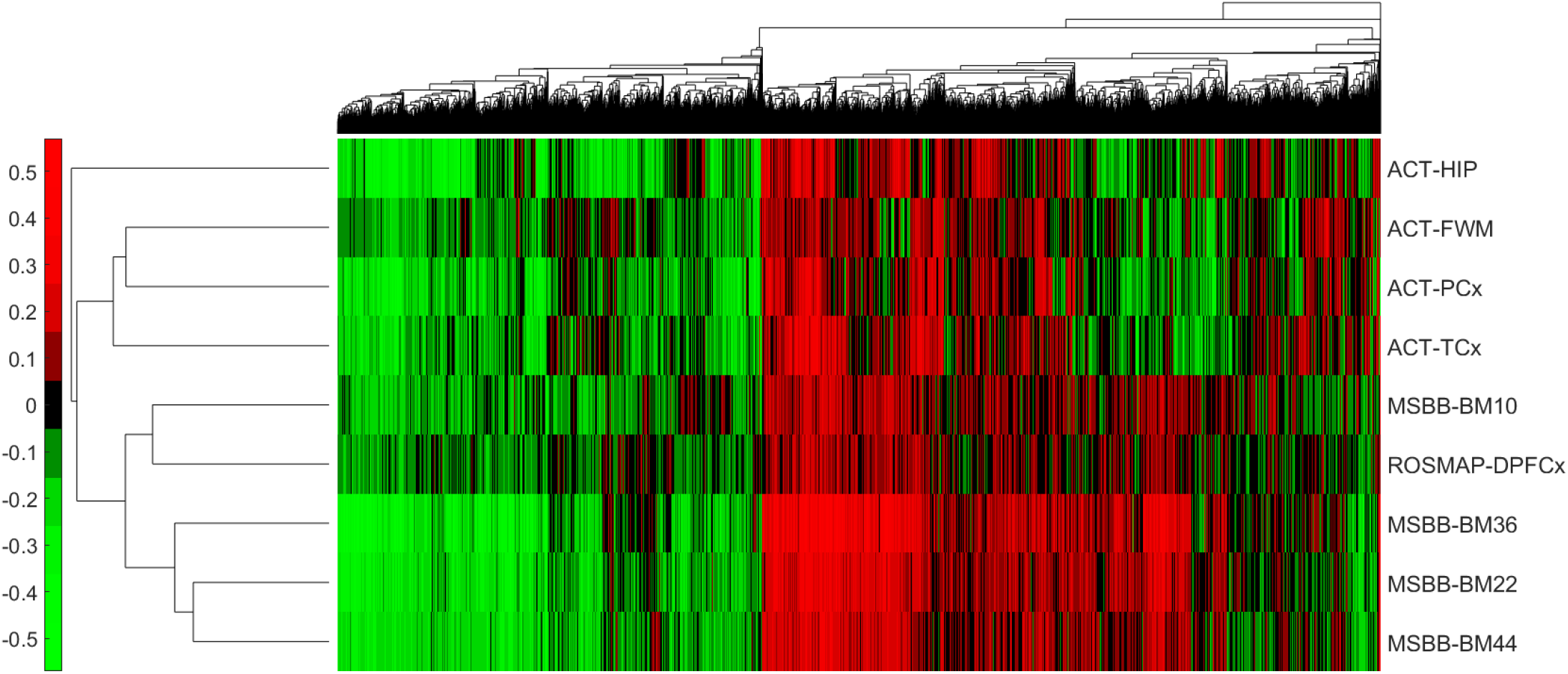
A heatmap of the associations between Aβ levels and gene expression across different brain regions. Each row corresponds to a region, and each column corresponds to a gene. Green indicates a negative association, and red a positive one.

### Result 2. Brain regions relevant to AD are highly weighted by DECODER

We hypothesize that the development of neuropathology is driven by similar molecular mechanisms in different brain regions, which is also supported by the results displayed in Table 2 and Figure 2. Therefore, focusing on the concordant molecular markers across different regions could lead to the discovery of reliable genes driving Aβ neuropathology. We propose a generative probabilistic model of neuropathology based on gene expression levels, and then we learn model parameters, each corresponding to a gene expression level’s true association with the neuropathology feature, concordant across brain regions (Methods). Starting with this probabilistic model, we learn the true expression-neuropathology associations in three different ways: (1) we equally weigh the observed associations in each brain region to maximize the data log-likelihood, (2) we combine brain regions with different weights so that this combination captures as much data variation as possible, and (3) we non-linearly combine individual region scores so that we can accurately reconstruct them from this combination (Methods). Each approach generates a single quantitative score for each gene representing its neuropathology marker potential, and we use these potentials to rank the genes. We refer to the new gene scores identified by these three approaches as the “equal weights score (EWS)”, “maximum variance score (MVS)”, and “best reconstruction score (BRS)”, respectively.

EWS and MVS both linearly combine regional associations (Methods). However, EWS equally weights the regions, while MVS learns the weights associated with different regions from the input data. We observed that, when applied to gene expression-Aβ associations, MVS highly weights regions previously known to be relevant to AD (Figure 3). The regions MSBB-BM36 and MSBB-BM22, which coincide with the medial temporal lobe (the brain part where AD-related cellular and structural alterations begin and have a more severe effect^22^), are the top two regions with the highest weights. White matter is suggested to be less vulnerable to AD than gray matter,^23^ and, interestingly, the white matter region ACT-FWM is assigned the lowest weight. Similarly, the dorsolateral prefrontal cortex region is affected in later stages of AD progression,^16^ and ROSMAP-DPFCx was also assigned a relatively lower weight.

**Figure 3.**
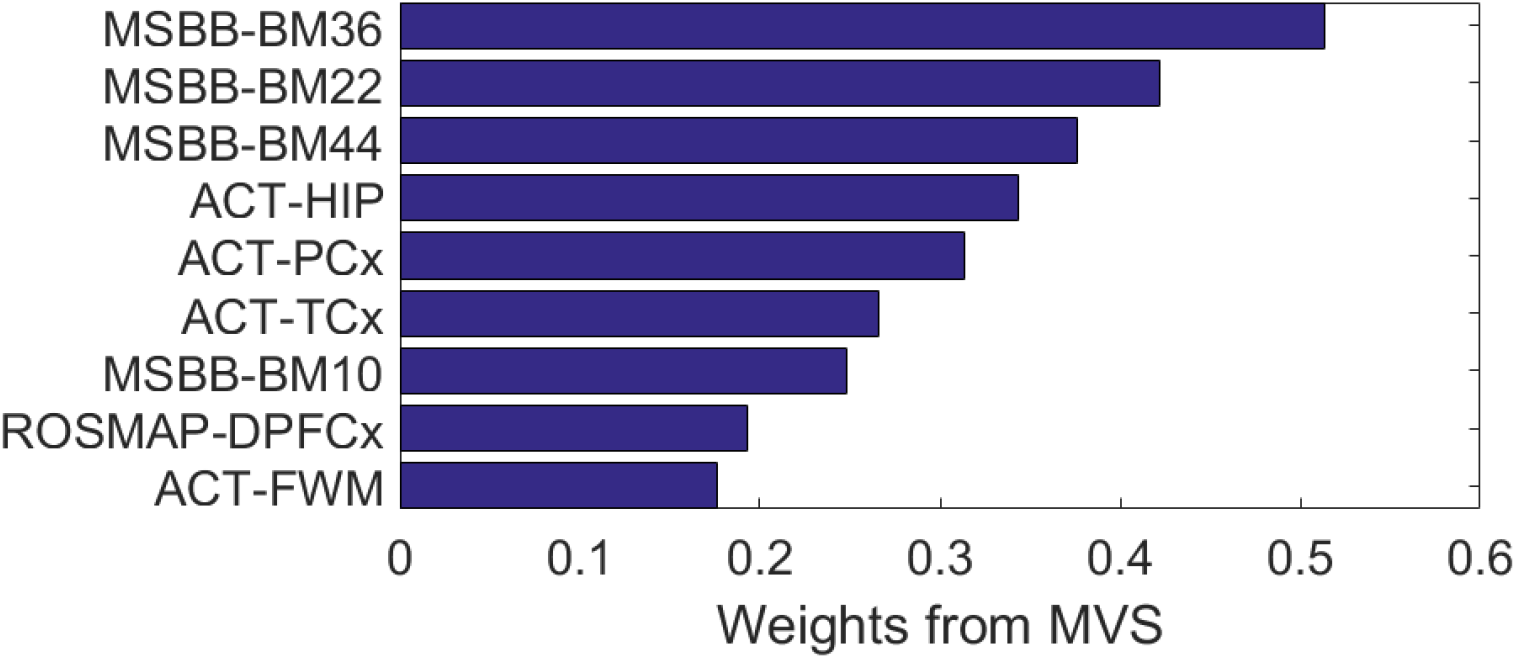
MVS weights associated with each of the nine brain regions for gene expression-Aβ associations. Brain regions are weighted consistently with their relevance to AD neuropathology.

BRS is an artificial neural network-based method that models latent space using a nonlinear combination of input variables. Thus, unlike EWS and MVS, BRS nonlinearly combines the individual regions (Methods). We conjecture that nonlinear embeddings could be useful for complex interactions among different regions that linear methods cannot capture. Scores computed by the three concordance-based approaches are highly consistent with each other, as shown in Supplementary Figure 1.

### Result 3. Concordance-based approaches identify statistically robust markers of AD pathology

We performed a cross-validation (CV) experiment to test whether the gene scores identified based on concordance across multiple regions (i.e., EWS, MVS, or BRS) are statistically more robust and informative than scores from individual region-based approaches (i.e., the association of gene expression with Aβ levels in each individual region). We also compared our concordance-based approaches to Fisher’s method, which is widely used to derive a single *p*-value from multiple associations with different significance levels (i.e., *p*-values). Fisher’s method assigns to each gene a score that is proportional to the mean negative logarithm of the individual *p*-values. In our experiments, we use Fisher’s method to combine nine *p* -values obtained by performing Pearson’s correlation test for each brain region.

In our CV experiment to test statistical consistency, for each of the nine regions used as a test fold, we used the other eight regions for training. For each test fold, we computed how well the gene scores from each of the eight individual regions, Fisher’s method, EWS, MVS, and BRS estimated test fold scores. Note that DECODER and Fisher’s method computed scores based on the eight training regions. We used a Spearman rank correlation between the actual vs. estimated gene scores to measure each method’s performance. We observed that our concordance-based approaches outperformed all alternatives in statistical consistency averaged across the nine regions used for testing (Figure 4). Thus, gene rankings are statistically more robust (i.e. likely to be replicated in an independent, unobserved dataset) when obtained based on the concordance of expression-neuropathology associations across regions compared to when determined based on an individual region or Fisher’s method. While the three concordance-based approaches performed similarly in terms of statistical consistency, we observed a very slight increase in performance with an increase in complexity of the approach we used to learn the scores (i.e., BRS performed slightly better than MVS, which in turn performed slightly better than EWS) (Methods).

**Figure 4.**
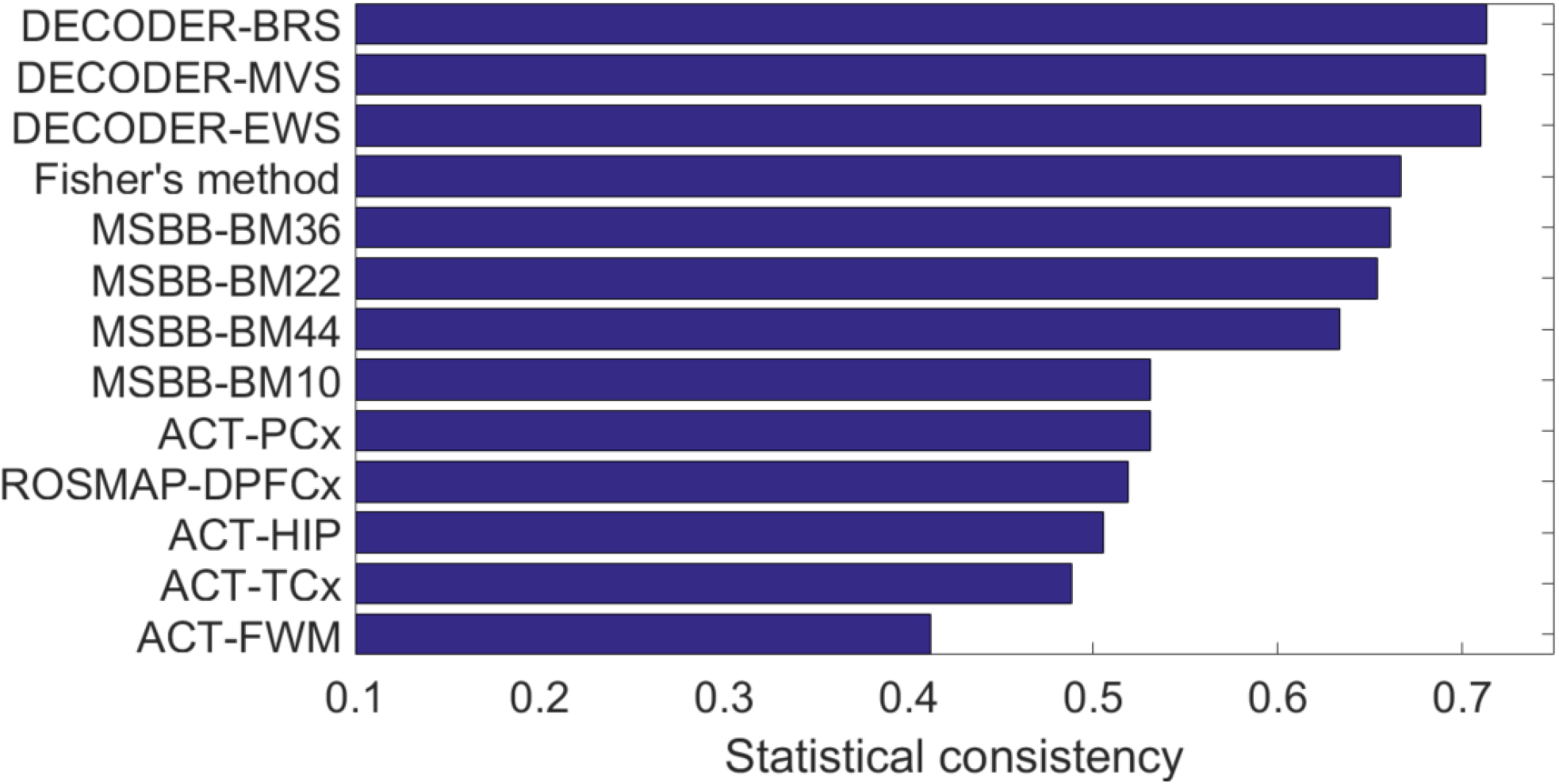
The statistical consistency achieved by DECODER, the commonly used Fisher’s method, and nine individual region scores for Aβ.

Fisher’s method works with *p*-values rather than observed associations. Therefore, to compute Fisher’s method consistency in a way comparable to other methods, we averaged consistency results over two different Fisher’s method choices: one choice combines *p*-values from a one-sided correlation test with an alternative hypothesis of negative correlation, and the other with an alternative hypothesis of positive correlation. Supplementary Figure 4 shows Fisher’s method bars separately from those two alternatives.

Fisher’s method achieves a relatively lower statistical consistency, likely because it generally puts a higher weight on highly-sampled regions since smaller association p-values are often obtained from those. However, highly sampled regions may not be the most relevant to AD. In our experimental setting of nine regions, ROSMAP-DPFCx includes the highest number of samples (Table 1). However, the dorsolateral prefrontal cortex is usually affected later as AD progresses^16^ and is therefore not as relevant to AD as several other regions with a smaller sample size in our data.

### Result 4. CEMs are highly enriched for genes known to be relevant to AD

In addition to statistical relevance, we also assessed the biological relevance of the top-scoring genes from the EWS, MVS, and BRS that we propose as part of our DECODER framework. To do this, we checked whether the CEMs identified by concordance-based approaches are more likely to be relevant to AD compared to the top genes identified by individual region-based approaches and Fisher’s method. The biological relevance metric we use is the number of genes overlapping with the ‘KEGG_ALZHEIMERS_DISEASE’ pathway (AD pathway) from the C2 collection (curated gene sets from online pathway databases) of the Molecular Signatures Database (MSigDB).^24^ This pathway contains 169 genes in total, 144 of which exist in all gene expression datasets from the nine brain regions we use in our experiments (Supplementary Table 1).

For each of the nine individual region-based approaches, Fisher’s method, EWS, MVS, and BRS, we selected the top N genes, where genes were sorted based on their score assigned by the method. Then, for each *n* = {1,…, *N*}, we analyzed the number of genes that overlap with the 144 AD pathway genes. We repeated the analysis for the “negative tail” and “positive tail” from each method; these terms refer to genes to which the method assigns highly negative scores and highly positive scores, respectively. Figure 5 presents the results averaged over the two tails for each method; the curves corresponding to EWS, MVS, and BRS (purple, green, and black solid curves, respectively) are above all other curves. This means that concordance-based approaches identify more true positives earlier than alternative approaches (i.e., individual region-based approaches represented by nine non-solid curves and Fisher’s method represented by the yellow solid curve). As mentioned earlier, one choice of Fisher’s method combines *p* -values from a one-sided correlation test with an alternative hypothesis of negative correlation, and the other with an alternative hypothesis of positive correlation. The negative and positive tails for Fisher’s method refer to the top genes from these alternatives, respectively.

**Figure 5.**
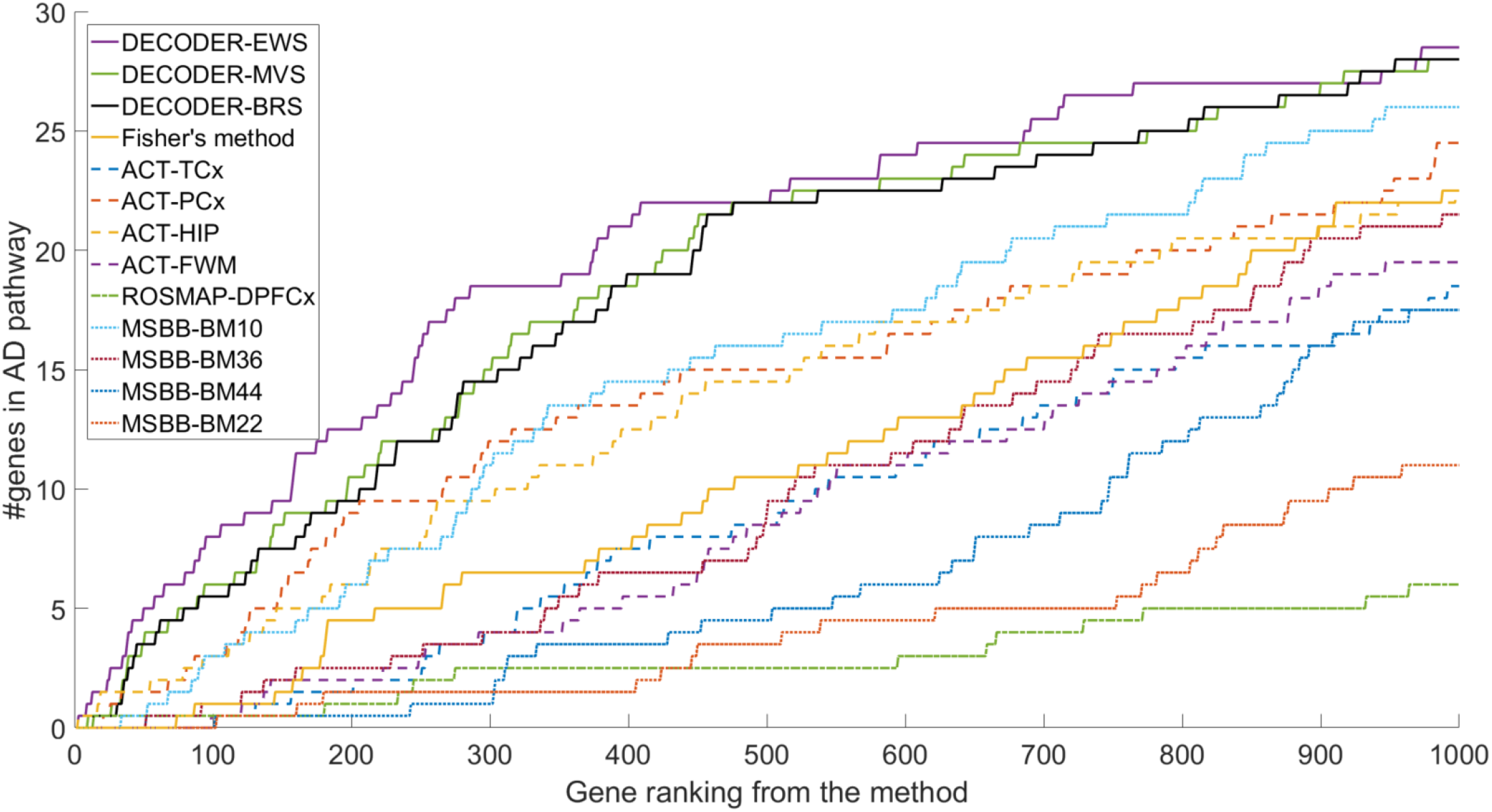
A comparison of the biological consistency achieved by three concordance-based scores we introduce (EWS, MVS, and BRS) to the commonly used Fisher’s method scores and nine individual region scores for Aβ. Consistency is measured by the overlap between the top n = {1,…, 1000} genes from each method (x-axis) and the 144 genes in the KEGG AD pathway.

Supplementary Figure 5 shows the comparative biological consistency results for up to 1,000 genes (*N* = 1,000) separately for the positive and negative tails for each method. Interestingly, the positive tail is not enriched for AD pathway genes; only a few of the top 1,000 positive tail CEMs are in the AD pathway (Supplementary Figure 5B), while several dozens of the top 1,000 negative tail CEMs are in the same pathway (Supplementary Figure 5A).

### Result 5. CEMs are highly enriched for pathways relevant to AD

To further investigate the biological relevance of CEMs, we examined the functional enrichment of 50 CEMs from the negative and positive tails of EWS, MVS, and BRS, where those tails correspond to genes that are assigned highly negative and highly positive scores, respectively. We considered 1,077 Reactome, BioCarta, and KEGG GeneSets (canonical pathways) from the C2 collection (curated gene sets from online pathway databases) of the MSigDB.^24^ We computed the significance of the overlap between each GeneSet and the 50 CEMs measured by Fisher’s exact test (FET) *p*-value and then applied false discovery rate (FDR) correction for multiple hypotheses testing for the 1,077 pathways.

The 50 CEMs from the Aβ negative tail of the EWS, MVS, and BRS were all significantly enriched for seven pathways (FET *p* ≤ 0.05), as shown in Table 3. These include known pathways for neurodegenerative diseases, including Parkinson’s disease and Huntington’s disease, in addition to AD. Other pathways in the table involved the electron transport chain or oxidative phosphorylation, implicated in a variety of neurodegenerative processes.^25–27^ Interestingly, the positive tail did not exhibit significant pathway enrichment for any of the three approaches. Since EWS generally achieved a more significant pathway enrichment than MVS or BRS (Table 3), we decided to pay particular attention to CEMs on the negative tail of EWS for our further experiments with CEMs.

**Table 3.** Fisher’s exact test (FET) p-values for the seven Reactome, BioCarta, and KEGG pathways for which the top 50 CEMs from the negative tail (i.e., top 50 genes with highly negative scores) of the EWS (2^nd^ column), MVS (3^rd^ column), or BRS (4^th^ column) were significantly enriched (p ≤ 0.05). All seven pathways contain the NDUFA9 gene.

| Pathway | FET $p$-value<br>from EWS | FET $p$-value<br>from MVS | FET $p$-value<br>from BRS |
| --- | --- | --- | --- |
| KEGG_OXIDATIVE_PHOSPHORYLATION | 9.0e-12 | 6.2e-7 | 6.2e-7 |
| REACTOME_RESPIRATORY_ELECTRON_TRANSPORT_ATP_S<br>YNTHESIS_BY_CHEMIOSMOTIC_COUPLING_AND_HEAT_PR<br>ODUCTION_BY_UNCOUPLING_PROTEINS_ | 9.0e-12 | 1.2e-5 | 1.2e-5 |
| KEGG_PARKINSONS_DISEASE | 1.9e-10 | 6.8e-6 | 6.8e-6 |
| REACTOME_TCA_CYCLE_AND_RESPIRATORY_ELECTRON_<br>TRANSPORT | 2.7e-10 | 8.9e-5 | 8.9e-5 |
| REACTOME_RESPIRATORY_ELECTRON_TRANSPORT | 9.3e-10 | 9.9e-4 | 9.9e-4 |
| KEGG_HUNTINGTONS_DISEASE | 3.5e-9 | 4.5e-5 | 4.5e-5 |
| KEGG_ALZHEIMERS_DISEASE | 2.9e-8 | 2.5e-4 | 2.5e-4 |

### Result 6. Concordance-based approaches are robust to different phenotypes/diseases, including AD tau phenotype and cancer survival

To test the applicability of the DECODER approach to other phenotypes or diseases, we also applied it to: (1) gene expression-tau associations from the nine brain regions (AD studies), and (2) gene expression-survival associations from 33 cancer types (The Cancer Genome Atlas (TCGA) study). Through these experiments, we aimed to obtain: (1) the gene markers whose expression levels are predictive of tau pathology across different brain regions, and (2) cancer patient survival across different cancers.

Figure 6 shows a heatmap of gene expression-tau neuropathology associations and the statistical and biological consistency results from applying DECODER and the alternative methods to these associations. Each row in the heatmap (Figure 6A) represents one of the nine brain regions in Table 1, and each column represents one of the 14,912 protein-coding genes whose expression was measured in all nine regions. As was the case in the heatmap for expression-Aβ associations (Figure 2), regions from the same study tend to group together in the heatmap for tau associations. Still, MSBB-BM44, a frontal cortex region from the MSBB study, connects to another frontal cortex region from a different study, ROSMAP-DPFCx, before it connects to other cortical regions from the MSBB study. Concordance-based approaches estimated unobserved test region scores more accurately than individual region-based approaches and almost as accurately as Fisher’s method (Figure 6B). Moreover, of 14,912 total genes, 144 genes in the ‘KEGG_ALZHEIMERS_DISEASE’ pathway (Supplementary Table 1) exhibited a large overlap with negative tail tau CEMs earlier than the highly ranked genes by individual region-based approaches or Fisher’s method (Figure 6C).

**Figure 6.**
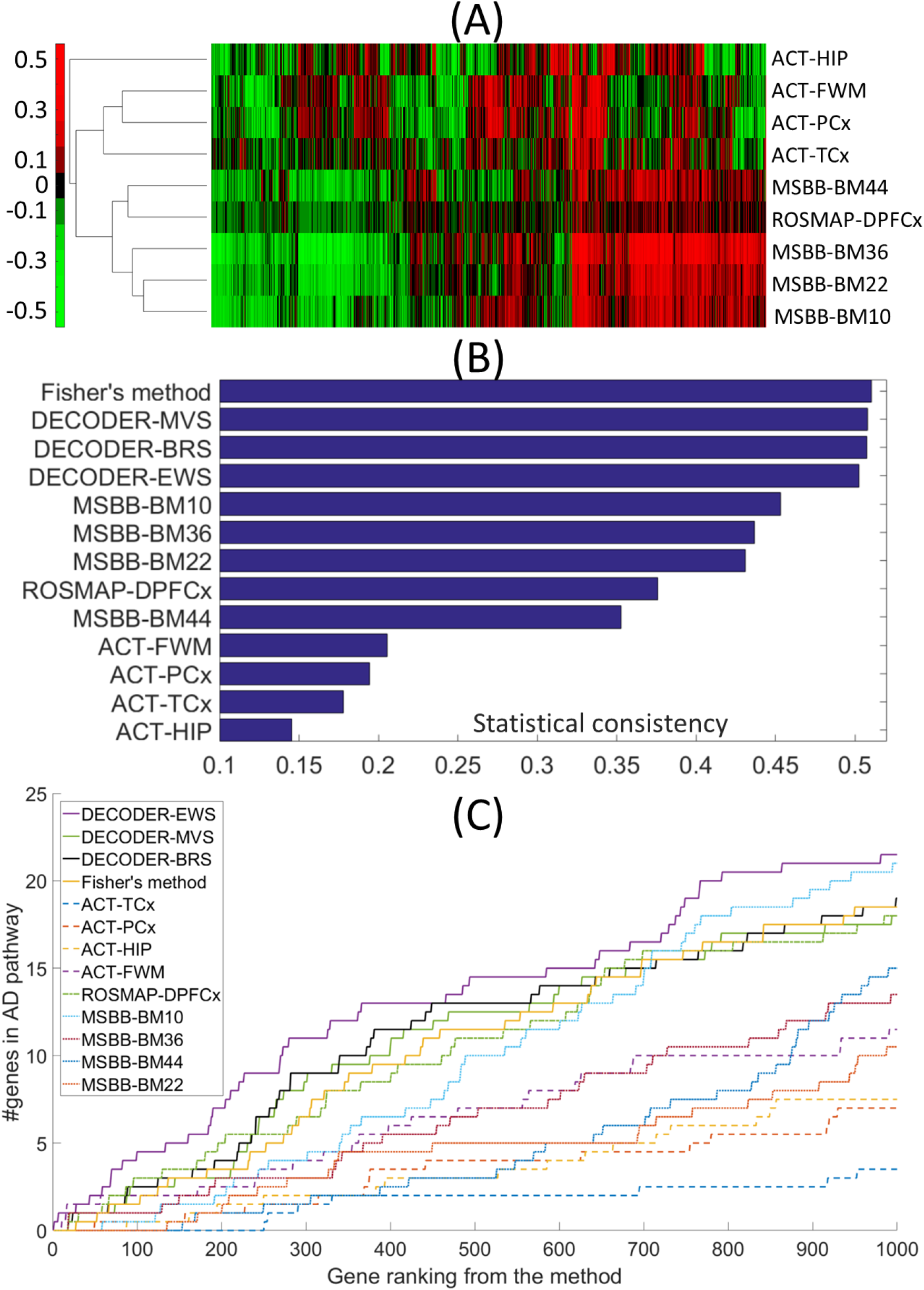
Testing of DECODER for gene expression-tau neuropathology associations. **(A)** A heatmap of the associations between tau levels and gene expression across different regions. **(B)** Comparison of concordance-based approaches to individual region-based approaches and Fisher’s method in terms of the estimation of unobserved test region scores. **(C)** Comparison of concordance-based approaches to individual region-based approaches and Fisher’s method in terms of overlap between KEGG AD pathway genes and the highly ranked genes by each approach.

Figure 7 shows a heatmap of TCGA gene expression-survival associations and the statistical and biological consistency results from applying DECODER and the alternative methods to these associations. Here, our hypothesis is that focusing on gene expression-survival associations concordant across different cancers may let us identify molecular markers generally important in cancer prognosis and may highlight prognosis-related molecular mechanisms in cancer.

**Figure 7.**
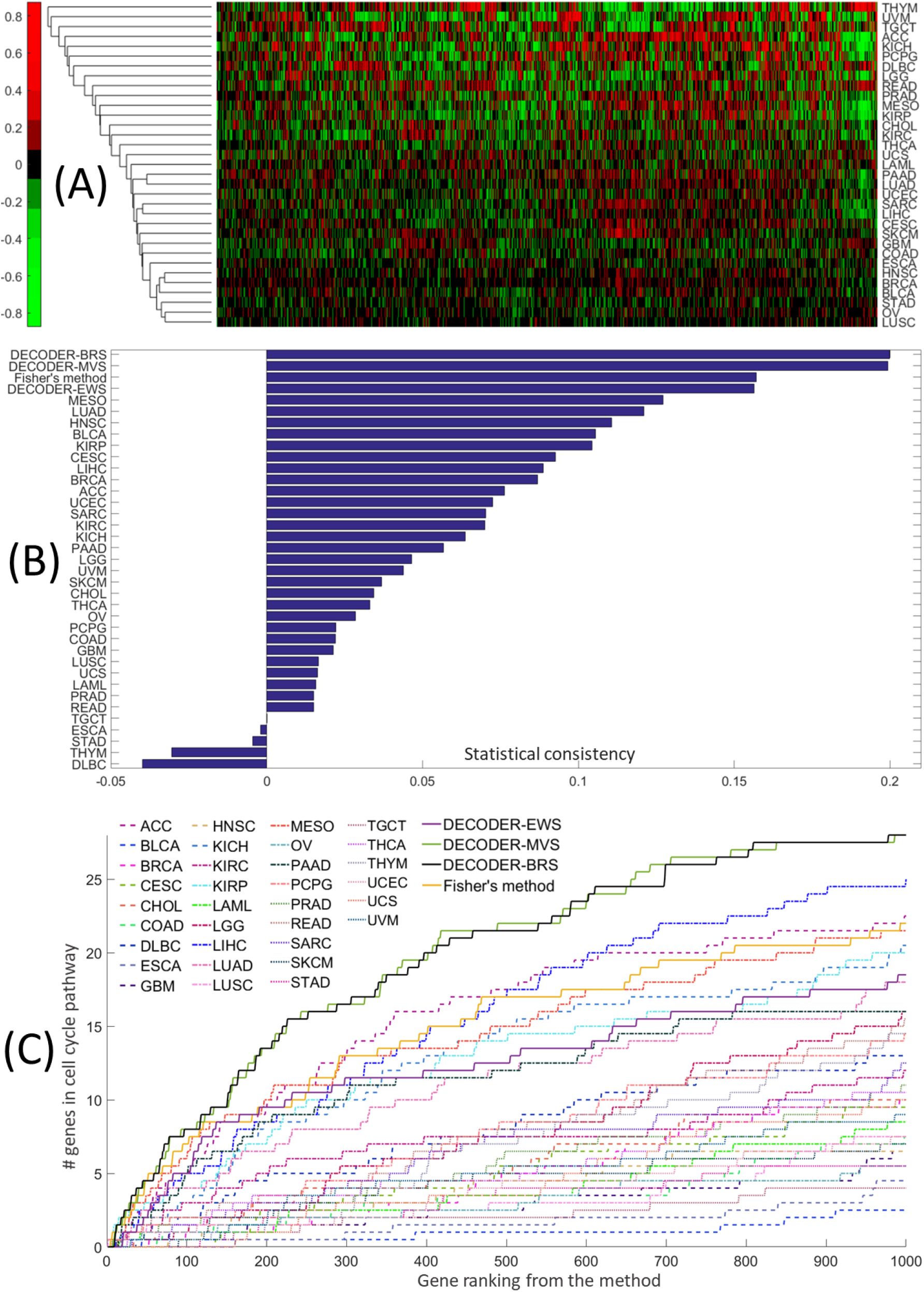
Testing of our concordance-based approaches for TCGA gene expression-survival associations from 33 cancer types. **(A)** A heatmap of the associations between survival and gene expression across different cancer types. **(B)** Comparison of concordance-based approaches to individual region-based approaches in terms of the estimation of unobserved test cancer type scores. **(C)** Comparison of concordance-based approaches to individual region-based approaches in terms of overlap between KEGG cell cycle pathway genes and the highly ranked genes by each approach.

Each row in the heatmap (Figure 7A) represents one of 33 cancer types, and each column represents one of the 15,097 genes included in the TCGA RNA-Seq data from all cancer types. Cancer is a heterogeneous disease, and survival is a phenotype that could be affected by a diverse set of factors. So, not surprisingly, we did not observe a visible clustering structure in the dendrogram of gene expression-phenotype associations. Still, concordance-based approaches, specifically MVS and BRS, estimated unobserved gene scores from the test cancer type (Figure 7B) more accurately than alternative approaches. Furthermore, of 15,097 total genes, 118 genes in the ‘KEGG_CELL_CYCLE’ pathway exhibited a large overlap with negative tail survival CEMs earlier than the highly ranked genes by individual region-based approaches or Fisher’s method (Figure 7C). This is not surprising given that the dysregulation of cell cycle-regulated genes is common in many cancers and has been associated with poor prognosis.^28–30^

Interestingly, we observe a clear outperformance of our relatively more complex approaches MVS and BRS over our simplest approach EWS in cancer survival marker identification problem, in terms of both statistical and biological consistency (Figure 7B-C). This might be because this problem poses a more challenging setting involving dozens of different tissue types and studies, and a more complex phenotype, survival, which is right-censored and highly affected by non-molecular factors.

### Result 7. The NDUFA9 gene is biologically validated in vivo to be a modifier of Aβ toxicity

To gain insight into the relevance of individual CEMs to Aβ toxicity, we utilized the nematode *C. elegans* as an animal model of Aβ proteotoxicity. Over the last two decades, *C. elegans* has been a particularly useful model system for studying genetic pathways that influence proteotoxicity since transgenic strains are easily constructed and maintained, and genes are efficiently knocked down by RNAi.^31,32^ We conducted experiments with a transgenic worm line displaying an age-associated aggregation of human Aβ_1-42_ peptide in their body wall muscle cells.^33^ The proteotoxic stress induced by the human transgene results in a rapid onset of age-associated paralysis.

To identify CEMs that function as modifiers of Aβ toxicity, we first identified nematode orthologs for the 20 CEMs on the negative tail from EWS (Table 4) using a stringent (BLAST *e*-value ≤ 10^-30^) reciprocal best hits (RBH) approach (Methods). This approach identified seven unique human genes with at least one RBH, highlighted in blue in Table 4. Interestingly, of those seven genes, only *NDUFA9*, a component of the mitochondrial electron transport chain (mETC) Complex I, was contained in any of the seven canonical pathways for which the top CEMs were enriched (Table 3), and *all* of these seven canonical pathways contained *NDUFA9*. For these reasons, we focused first on the worm ortholog of *NDUFA9*, encoded by the gene Y53G8AL.2.

**Table 4.**
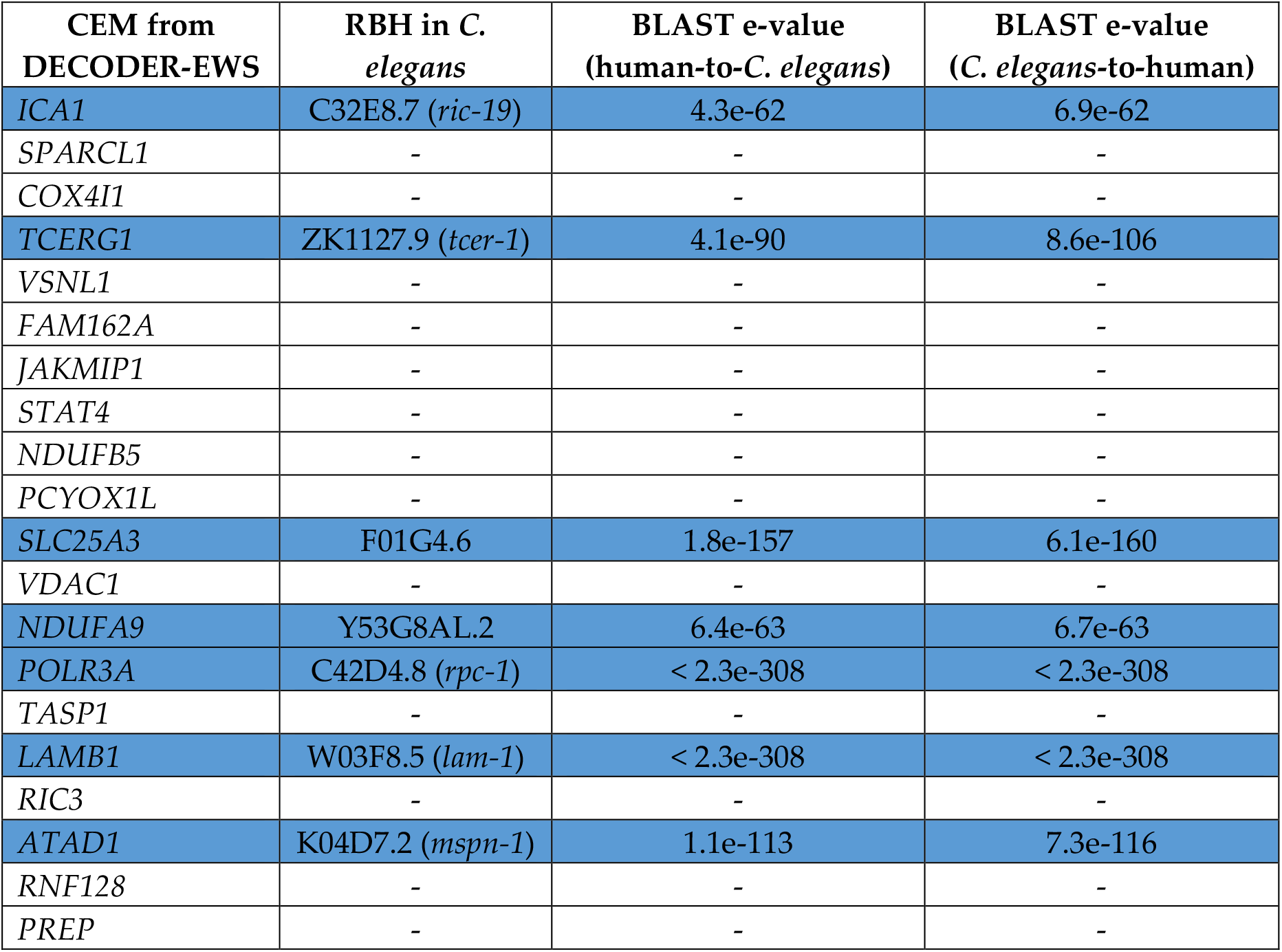
The 20 negative tail CEMs from EWS (1^st^ column) sorted by their absolute scores, reciprocal best hits (RBHs) of the gene in C. elegans (2^nd^ column), BLAST e-values of human-to-C. elegans, and C. elegans-to-human mapping (3^rd^ and 4^th^ columns, respectively). The rows corresponding to the CEMs that have an RBH in C. elegans are highlighted in blue.

To test candidate modifiers of Aβ1-42 toxicity, we used RNAi by bacterial feeding to reduce expression of the selected target gene.^34^ Worms expressing the Aβ_1-42_ transgene fed on bacteria expressing Y53G8AL.2 RNAi displayed a significant delay in paralysis compared to isogenic animals fed on bacteria expressing an identical RNAi vector that does not contain a dsRNA-coding region — empty vector (EV) — (Figure 8A). Suppression of paralysis by knockdown of Y53G8AL.2 was comparable to RNAi knockdown of the insulin-like receptor gene *daf-2* (Figure 8B), one of the strongest known suppressors of Aβ toxicity in worms^35^.

**Figure 8.**
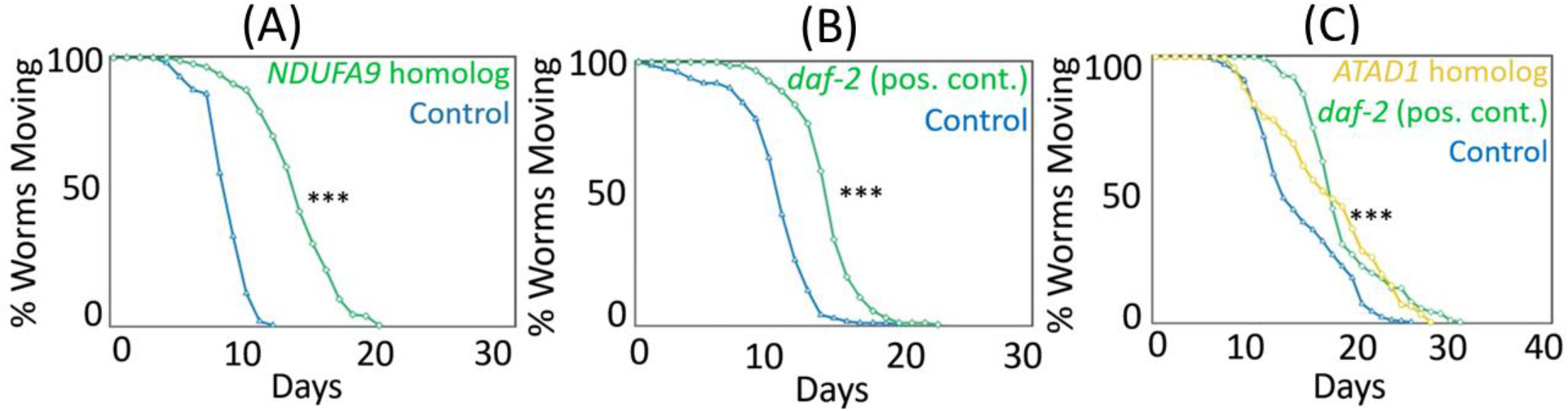
The worm NDUFA9 homolog promotes Aβ toxicity in C. elegans. **(A)** RNAi knockdown of the NDUFA9 homolog Y53G8AL.2 strongly suppresses Aβ-induced paralysis. **(B)** Y53G8AL.2’s suppression of paralysis is comparable to the potent proteotoxicity suppression treatment with daf-2 RNAi. **(C)** RNAi knockdown of the C. elegans homolog for ATAD1, another CEM from the negative tail of the Aβ EWS ranking that is relevant to ATP metabolism, also significantly suppresses Aβ-induced paralysis. *** p-value less than 1e-3.

Y53G8AL.2 and *NDUFA9* have a strong sequence conservation (63% protein sequence). In humans, *NDUFA9* is the alpha subcomplex subunit 9 of the enzyme complex Complex I, also known as NADH (Nicotinamide Adenine Dinucleotide Hydrate) dehydrogenase or NADH:ubiquinone oxidoreductase. Complex I, the first and largest enzyme complex in the mETC, is located in the inner mitochondrial membrane and catalyzes electron transfer from NADH to ubiquinone.^36^

We note that identifying the *NDUFA9* gene would not be possible by examining the top genes from a single region or Fisher’s method, as shown in Table 5. Concordance-based scores led to a high ranking of the *NDUFA9* gene and made it possible to computationally identify its significance, which led to its biological validation in *C. elegans*.

**Table 5.** NDUFA9 ranking on the negative tail for Aβ associations based on concordance-based scores, Fisher’s method, and individual region-based scores.

| Method | *NDUFA9* ranking |
| --- | --- |
| DECODER-EWS | 13 |
| DECODER-MVS | 49 |
| DECODER-BRS | 45 |
| Fisher's method | 178 |
| ROSMAP-DPFCx | 234 |
| ACT-TCx | 320 |
| ACT-PCx | 182 |
| ACT-HIP | 250 |

| Method | NDUFA9 ranking |
| --- | --- |
| ACT-FWM | 433 |
| MSBB-BM10 | 623 |
| MSBB-BM36 | 874 |
| MSBB-BM44 | 1177 |
| MSBB-BM22 | 1064 |

Given that NDUFA9 — a gene with an important role in ATP synthesis and mitochondrial function — significantly suppressed paralysis, we also performed RNAi knockdown experiments to test the C. elegans homolog of *ATAD1*, the other CEM in Table 4 that is relevant to the ATP metabolism. Prior research showed that *ATAD1* maintained mitochondrial function.^37^ *ATAD1* is a member of the ATPase family which is a class of enzymes that catalyze the decomposition of ATP into ADP. ATPase plays a critical role in regulating the surface expression of AMPA (α-amino-3-hydroxy-5-methyl-4-isoxazolepropionic acid) receptors that are ionotropic transmembrane receptors for glutamate and mediate fast synaptic transmission in the central nervous system. AMPA receptor-related changes are a core feature of age-related cognitive decline.^38^ Thus, *ATAD1* plays an important role in regulating synaptic plasticity as well as learning and memory. We observed a significant suppression of paralysis in the *C. elegans* Aβ strain when it was treated with *ATAD1* homolog K04D7.2 *(mspn-1)* (Figure 8C).

### Result 8. Knockdown of NDUFA9’s nematode homolog strongly decreases whole animal oxygen consumption

Y53G8AL.2, the nematode homolog of the human *NDUFA9* gene (Table 4), showed robust suppression of paralysis in our RNAi knockdown experiments in *C. elegans* (Figure 8A). Consistent with its homology to human *NDUFA9* — an important component of the mETC — Y53G8AL.2 has been shown to impact oxidative phosphorylation in isolated mitochondria.^39^ However, no studies have directly investigated the impact of Y53G8AL.2 on organismal respiration. Because Y53G8AL.2 has not been extensively studied in *C. elegans*, we sought to confirm that the gene encodes a functional component of the mETC, as predicted by its homology to *NDUFA9*. Consistent with such a role, we observed that RNAi knockdown of Y53G8AL.2 strongly decreases whole animal oxygen consumption (Methods) (Figure 9).

**Figure 9.**
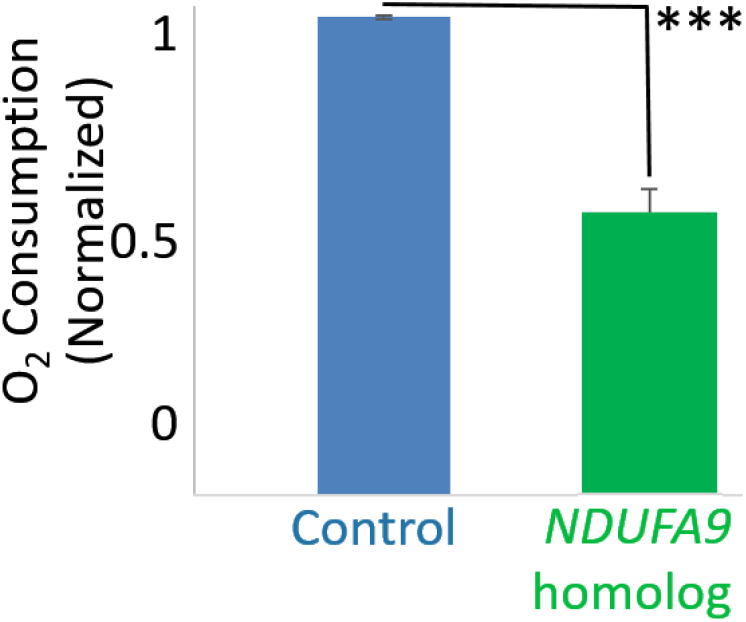
Knockdown of NDUFA9 homolog Y53G8AL.2 greatly reduces whole animal oxygen consumption. O2 consumption is normalized to the O2 consumption rate of one animal on control RNAi (EV). *** p-value less than 1e-3.

Thus, we concluded that reduced expression of *NDUFA9* homolog Y53G8AL.2 impairs the function of Complex I of the mETC, which robustly attenuates Aβ toxicity (Figure 8A).

### Result 9. In vivo validation of 12 additional Complex I genes identifies mitochondrial Complex I as a potential AD drug target

Thus far, we have established that: (1) CEMs were significantly enriched for functional categories relevant to mitochondrial respiration, ATP synthesis, and mETC, all including *NDUFA9* (Table 3), and (2) knockdown of the nematode homolog of *NDUFA9*, a subunit of mETC Complex I, strongly suppressed paralysis (Figure 8A) and significantly decreased whole animal oxygen consumption in *C. elegans* (Figure 9). We therefore hypothesized that Complex I may act as a key regulator of Aβ toxicity. We first sought to confirm that the Complex I genes exhibit a cluster structure in the 2-dimensional space that explains the highest amount of variance in the gene expression-Aβ associations from nine regions (i.e., the space of the first two principal components). Supplementary Figure 2 shows that most of the Complex I genes, represented by blue dots, are projected very closely to each other in the first two principal components space of expression-Aβ associations. This observation suggests that Complex I genes have similar Aβ association levels, and that other Complex I genes might also function, like Y53G8AL.2, as modifiers of Aβ-induced toxicity. To test this hypothesis, we individually knocked down 12 additional *C. elegans* genes that encode homologs of human Complex I proteins (Table 6) in the Aβ_1-42_ worms. Strikingly, knockdown of any of these 12 additional Complex I RNAi clones significantly delayed paralysis, with several exceeding the effect of Y53G8AL.2 (Figure 10). These computational and biological findings pinpoint Complex I of the mETC as a critical mediator of Aβ proteostasis.

**Table 6.**
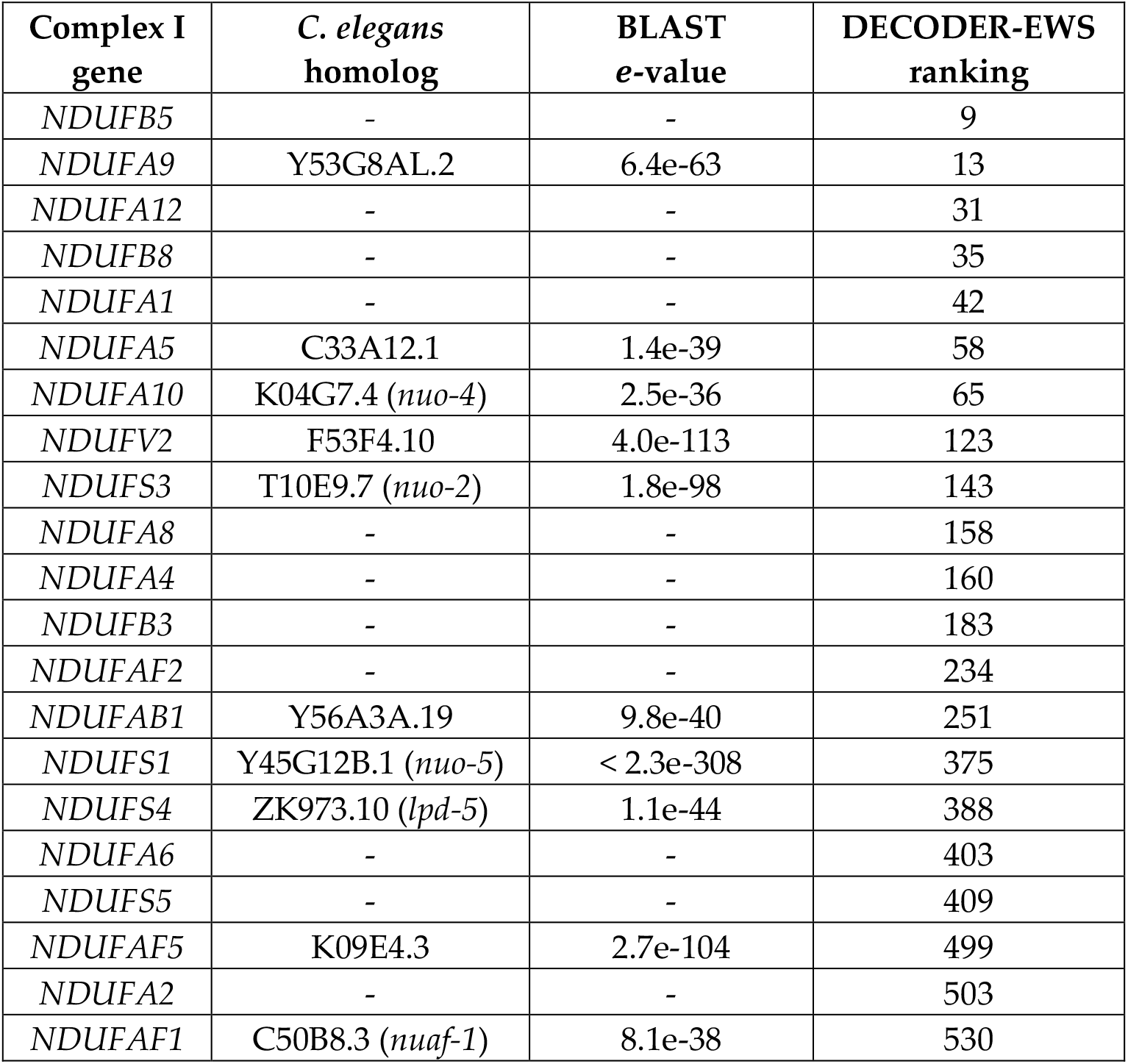
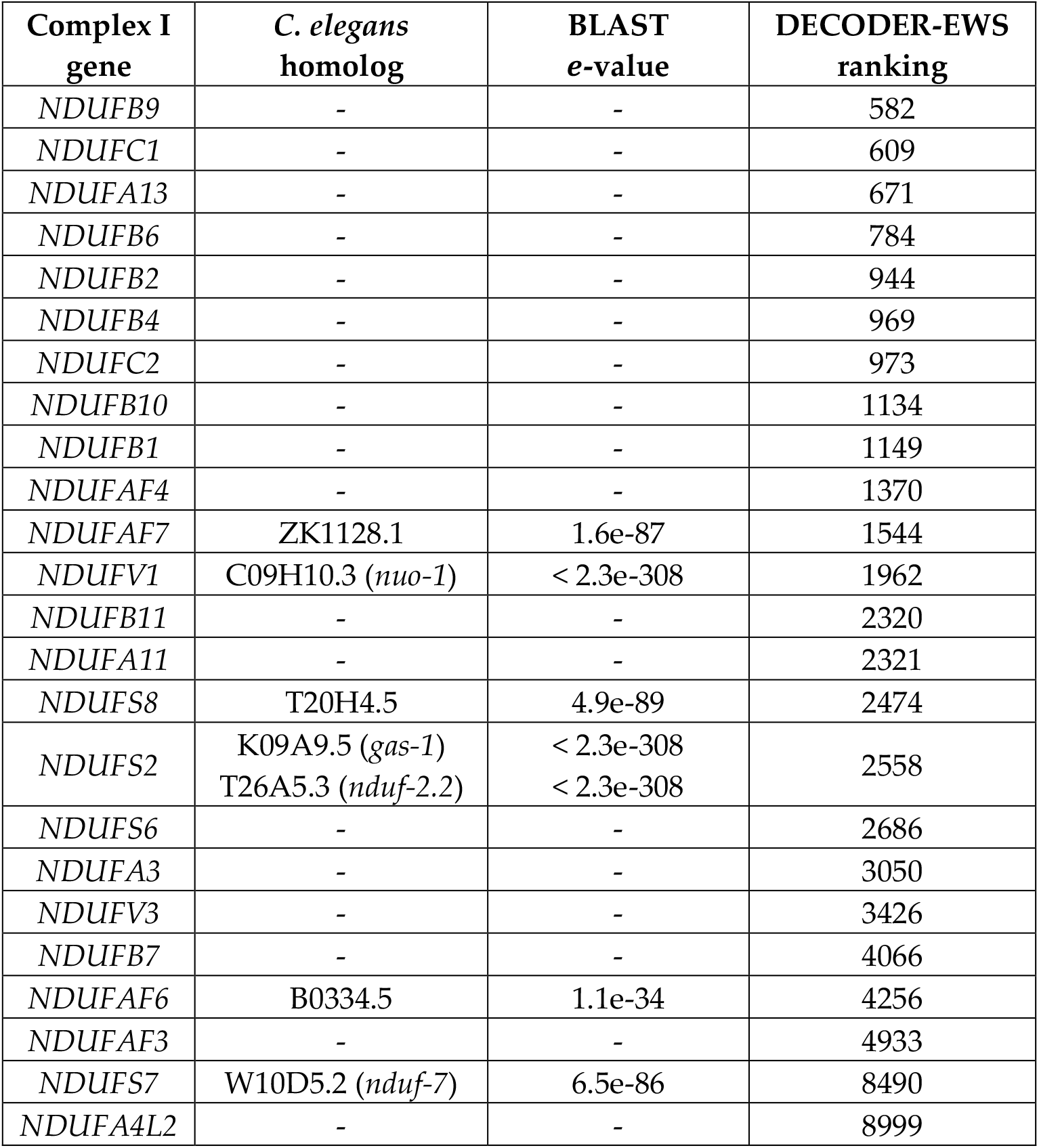
Human Complex I genes (1^st^ column), the strong ortholog of the NDUF gene in C. elegans (2^nd^ column), BLAST e-values of human-to-C. elegans mapping (3^rd^ column), and the Aβ negative tail ranking of the gene from our concordance-based EWS method (4^th^ column).

**Figure 10.**
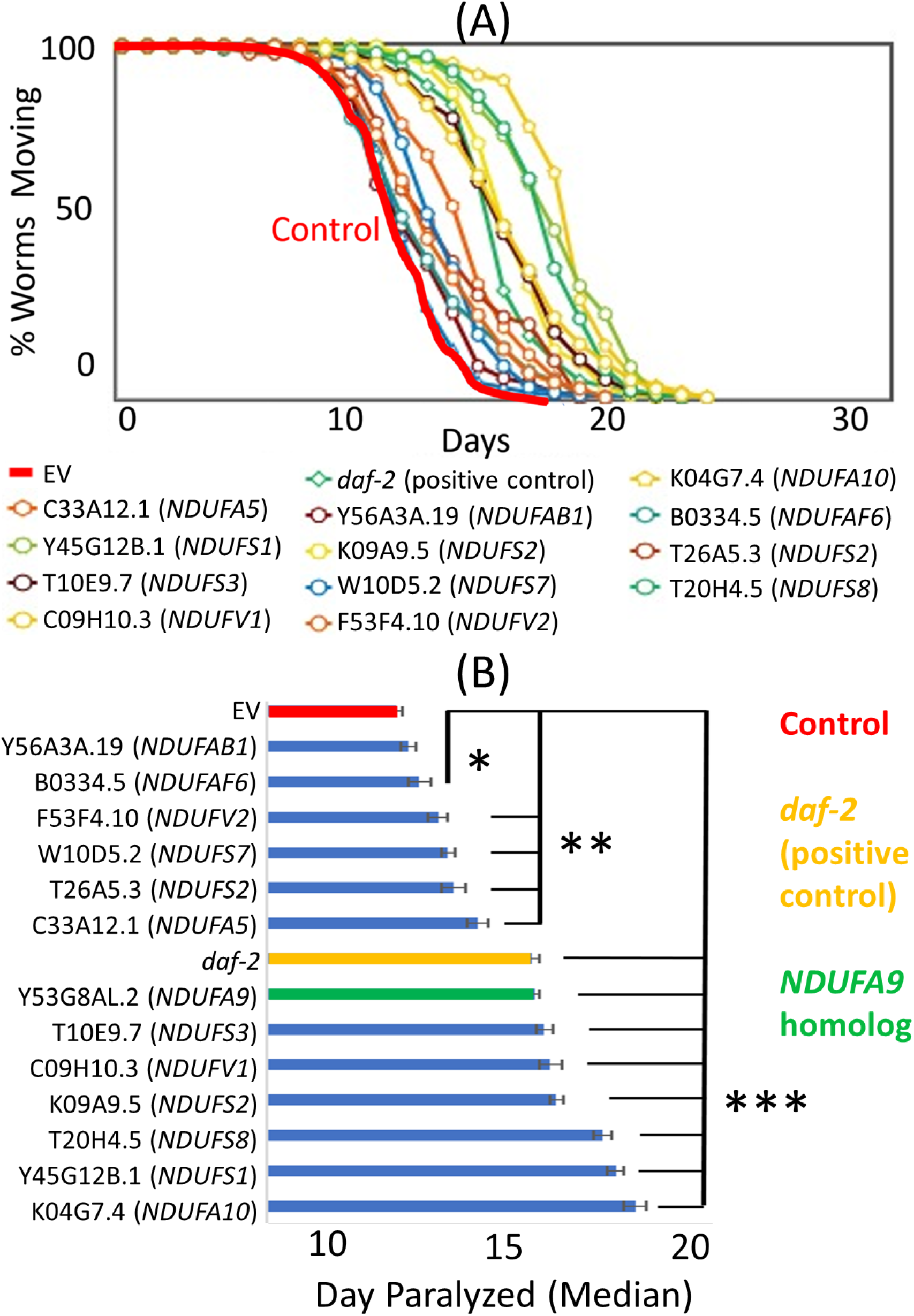
Knockdown of mitochondrial Complex I genes’ homologs in C. elegans robustly protects against Aβ toxicity. **(A)** Paralysis curves for the reciprocal best orthologs of human Complex I genes. All tested RNAi conditions significantly suppressed paralysis. **(B)** The same data as in (A) plotted as the median day of paralysis for each population of worms. Half of the conditions showed even stronger suppression than daf-2 and Y53GAL.2 RNAi conditions. Error bars mean standard error of the mean across experiments. * p-value < 0.05. ** p-value < 0.01. *** p-value < 0.001.

### Result 10. A multi-omic module subnetwork involving Complex I highlights mechanisms relevant to AD

Given our conclusion that mETC Complex I is a critical mediator of Aβ proteostasis, another important question concerns how Complex I is regulated in this process. Molecular processes are orchestrated by a complex interplay between different genetic and epigenetic regulatory elements. The ROSMAP study^12,13^ provides DNA methylation and microRNA (miRNA) measurements from the same individuals in addition to mRNA expression data for 542 individuals used in our concordance-based gene ranking framework (Table 1). These additional data types let us investigate how Complex I is regulated in a multi-omic setting.

We applied the MGL^40^ algorithm to a data matrix of 35,354 variables from three types of molecular data – mRNA, DNA methylation, and miRNA — in the ROSMAP study and learned a multi-omic module network that might shed light on the regulatory mechanisms in AD (Methods). DNA methylation and miRNAs are important epigenetic mechanisms posited to play an important role in AD, especially because environmental factors affect aging through epigenetic modifications in an individual.^41^

Figure 11 shows the subnetwork of the learned module network that represents the modular dependency structure for the Complex I modules. This subnetwork includes ten modules that are: (1) connected to at least three of the eight modules that are significantly enriched (FET *p* ≤ 0.05) for Complex I genes, and (2) either significantly enriched for a functional pathway or contain either a DNA methylation site or a miRNA. Next to each module in Figure 11, we list the top pathway for which this module is significantly enriched. The four orange-highlighted modules are significantly enriched for Complex I genes. We note that although not at the top in their enriched pathway list, ‘KEGG_ALZHEIMERS_DISEASE’ is a pathway for which each of these four modules is significantly enriched. See Supplementary Table 3 for detailed information on all 500 modules learned by the MGL algorithm (Methods).

**Figure 11.**
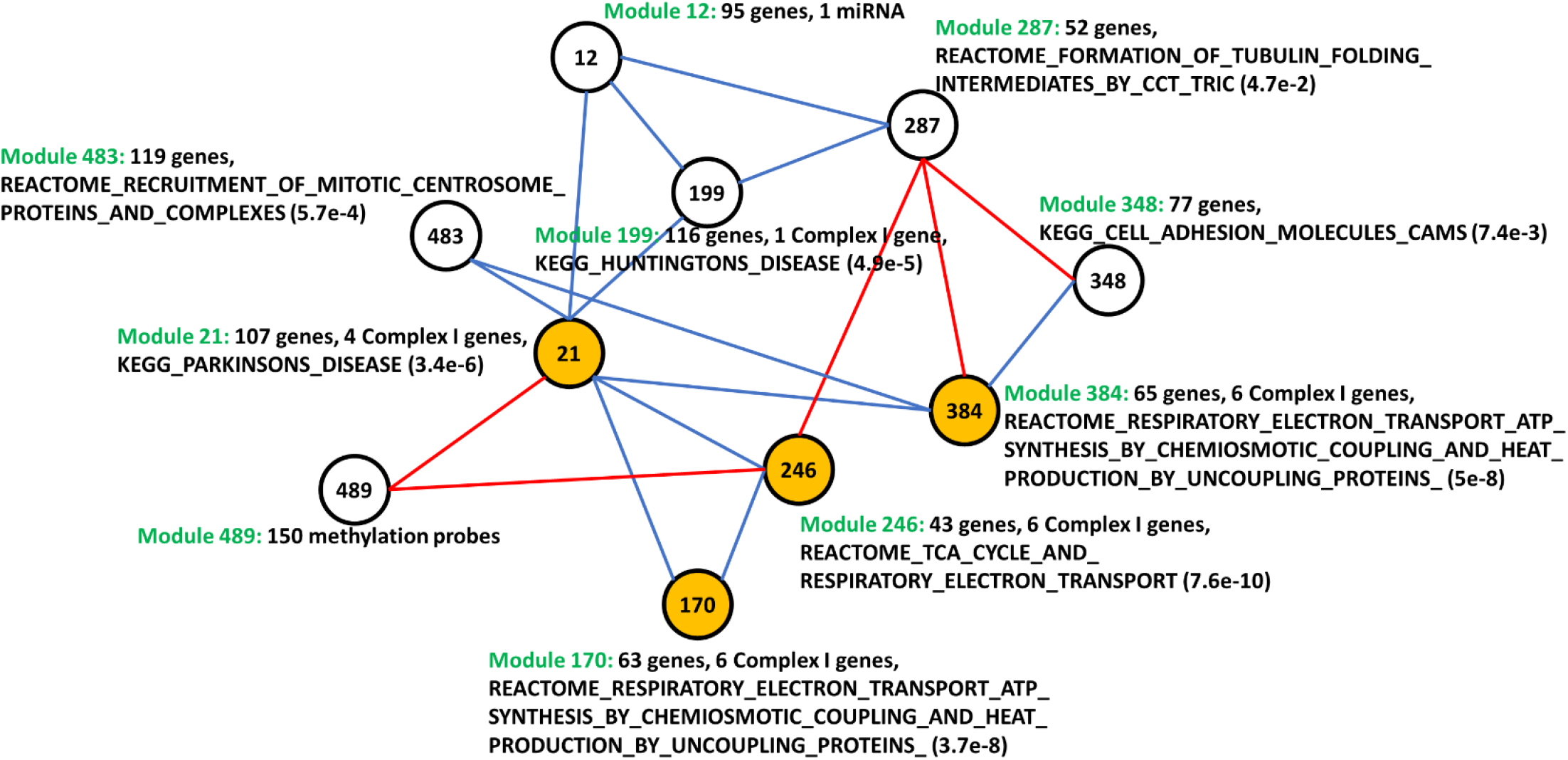
A multi-omic subnetwork of Complex I, learned by the MGL algorithm. Module 489 contains 150 methylation sites, and module 12 contains one miRNA (hsa-miR-633); other modules contain only mRNA variables. Blue edges represent up-regulation, and red edges represent down-regulation, between two connected modules.

Eight of the ten modules in the Figure 11 subnetwork contain only genes (mRNAs), and each of these modules is significantly enriched for pathways that are relevant to AD. The other two modules — modules 12 and 489 — contain non-mRNA variables (i.e., methylation or miRNA), and they are both connected to module 21, which is the hub in the subnetwork (i.e., connected to many other modules) and significantly enriched for Complex I genes. It was suggested earlier that reduction in mitochondrial function alters the expression of genes that mediate DNA methylation.^42^ It was also suggested that increased Aβ levels promote the production of reactive oxygen species that cause DNA oxidation; and DNA methylation events modify susceptibility to DNA oxidation, resulting in the acceleration of neurodegenerative events.^43^ Therefore, it is not surprising that module 21 has a dependence on module 489, which contains 150 methylation sites. Supplementary Table 2 lists the 80 unique genes to which these 150 methylation sites are mapped. The other module connected to module 21 and containing non-mRNA variables, module 12, contains miRNA hsa-miR-633, which was shown to be down-regulated in AD.^44^ Given that *NDUFA9* and most other Complex I genes were highly ranked on the negative tail of concordance-based gene scores for Aβ (i.e., down-regulated in AD) (Supplementary Table 4), an up-regulation between module 21 and module 12 is not surprising. This up-regulation is represented by the blue edge between modules 21 and 12 in Figure 11. We observed that the 95 genes in module 12 significantly overlap (*p* < 0.05) with the 1,231 target genes of hsa-miR-633 that are predicted by miRDB,^45^ which employs MirTarget^46^ algorithm version 3.0.

## Discussion

Alzheimer’s disease (AD) is a progressive neurodegenerative disorder with no cure. The molecular mechanisms underlying AD neuropathology remain unknown, and it is critical to identify true positive biomarkers for these mechanisms. An increasing number of AD studies provide different types of brain tissue data, including gene expression and neuropathological phenotypes. This offers a unique opportunity to perform coordinated analyses leveraging the different samples and data types now available.

Earlier research efforts have successfully reduced false positives by focusing on concordant associations across different molecular data types from the same study. White et al.^47^ and Yang et al.^48^ analyzed ROSMAP data, examining single nucleotide polymorphisms (SNPs) with significant phenotype association p-values only if their methylation status and/or mRNA levels were also associated with the phenotype. An important prior study that performed multi-regional or multi-study analyses of AD gene expression data was done by Wang et al.^14^, who employed an integrative network analysis approach using MSBB microarray data from 19 brain regions. Examining the gene expression similarities between regions, the authors detected strong interactions across regions with strong physical interconnectivity. Zhang et al.^10^ generated gene regulatory networks from three brain regions using an integrated systems approach and examined the functional network organization and its relation to AD pathology. However, both studies compared findings across different regions; they did not analyze the concordance of pathological molecular mechanisms across regions. Xu et al.^49^ compiled publicly available AD expression data from four brain regions measured in a total of 1,246 samples in 20 datasets. They identified genes that are differentially expressed between AD and control samples in all four regions and prioritized potential regulators of those genes by integrating data from GWAS, brain expression quantitative trait loci (eQTL), protein-protein interaction (PPI), and AD mouse models.

In this work, we introduced DECODER, which probabilistically models observed data to capture concordant associations between gene expression and neuropathology across different brain regions. We learned model parameters using three different approaches —equal weights score (EWS), maximum variance score (MVS), and best reconstruction score (BRS). Each approach assigned a score to each gene, enabling us to rank genes based on the level of agreement of brain regions on the association between gene expression and neuropathology. We demonstrated that each concordance-based approach reduces the rate of false positive associations and, thus, increases the chance that the identified genes are relevant to AD. This process led us to a short list of potential AD neuropathology biomarker genes, which we call concordant expression markers (CEMs), that are highly ranked by our concordance-based approaches. CEMs were enriched for several pathways relevant to the mitochondrial electron transport chain (mETC). The gene *NDUFA9*, involved in mETC, is a highly ranked CEM. This gene was identified to be differentially down-regulated between cognitively normal controls and AD individuals in four brain regions and significantly correlated with Aβ in mouse models.^49^ We tested *NDUFA9* and 12 other Complex I genes in *C. elegans* Aβ proteotoxicity models and showed that knockdown of each Complex I gene significantly suppresses Aβ toxicity.

Our findings suggest that mild Complex I inhibition could be a new paradigm for developing future AD therapeutics. Although it may seem paradoxical that mild inhibition of Complex I is protective, given that Complex I expression is lower in the brains of AD patients compared to healthy controls, a reduction in gene expression could simply reflect a protective response to pathological changes occurring during the disease. Indeed, several recent studies have indicated that mild mitochondrial dysfunction can induce a variety of protective responses including the mitochondrial unfolded protein response, mitophagy, and cytosolic chaperones. These protective responses to mitochondrial stress have been collectively referred to as mitohormesis.^50,51^ Additional support for Complex I as a particularly effective target can be found from prior work has indicated that mild inhibition of Complex I with the small molecule CP2 reduced Aβ and tau levels in mouse models of familial AD.^52^ Capsaicin is a natural product that can inhibit Complex I, and a capsaicin-rich diet was associated with lower total serum Aβ levels in the elderly.^53^ Furthermore, capsaicin reduced AD-associated tau changes in the hippocampus of type 2 diabetes rats.^54^ These earlier studies’ findings, combined with our novel results from the brain region concordance-based approaches, suggest that Complex I is indeed a promising potential pharmacological avenue toward treating AD.

The *C. elegans* gene *K04G7.4* (*nuo-4*) — ortholog of the human Complex I gene *NDUFA10* — was the strongest suppressor of Aβ toxicity, a gene shown to be significantly enriched in serotonergic neurons compared to whole-animal expression.^55^ Given that our *C. elegans* model expresses human Aβ_1-42_ peptide solely in body wall muscle cells, this suggests the possibility that cell non-autonomous, neuron-to-muscle signaling mediates the regulation of Aβ toxicity following Complex I inhibition in *C. elegans*. Prior studies using *C. elegans* have shown that neuronal signaling and endocrine pathways are integrated into the physiology of other tissues in the organism, including the regulation of heat shock response modulation of protein homeostasis in post-synaptic muscle cells.^56–58^ Interestingly, neuronal RNAi of mETC genes is sufficient to extend lifespan in worms and can induce changes in gene expression in the intestine including activation of the mitochondrial unfolded protein response.^59^ This mitochondrial stress response has been proposed to increase lifespan^59^ and recently implicated in suppression of Aβ toxicity in worms^60^. However, other studies have shown that the lifespan extension is independent of the mitochondrial unfolded protein response.^61–64^ It will be of interest to determine whether activation of the mitochondrial unfolded protein response plays a role in suppression of Aβ toxicity following Complex I inhibition, and whether these effects of Complex I inhibition are cell-autonomous or cell non-autonomous.

One limitation of our computational experiments to test DECODER’s performance on identifying neuropathology marker genes is that the neuropathological phenotype quantifications we used for some of the nine brain regions had not been obtained exactly from that phenotype or region. For the MSBB RNA-Seq samples, no tau, tangle, or Aβ quantifications were provided. Therefore, we used Braak stages as a proxy for tau levels since Braak staging is based on the regional distribution pattern of neurofibrillary tangle density across the brain,^65^ and we used the neuritic plaque density mean across five cortical regions as a proxy for Aβ. Thus, the samples from the four different MSBB regions were assigned the same values for Aβ and the same values for tau. Moreover, histelide^66^ Aβ quantification was provided for only two of the four ACT regions — ACT-TCx and ACT-PCx. Therefore, we used ACT-PCx Aβ levels for ACT-FWM and ACT-TCx Aβ levels for ACT-HIP based on regional proximity. We surmise that regions from the same study tend to group together in the heatmaps in Figure 2 and Figure 6 partly because of these study-specific arrangements we made due to limited data availability.

DECODER is a general framework applicable to any omic data type (e.g., proteomic, metabolomic) besides transcriptomic data as long as it is possible to get association statistics, such as correlation coefficients, from the data. Therefore, we expect a wider availability of AD multi-omic data to let us apply DECODER to identify concordant neuropathology markers from other molecular data types as well.

Another important direction for future work is to develop a systematic approach to iterate between a computational algorithm phase and a biological validation phase. At each iteration, the computational algorithm can be improved using causal markers identified by the experiments in the previous biological validation phase, which could lead to a new set of genes to test in *C. elegans* and/or other model organisms in the next biological validation phase. Biological validation experiments let us identify causal relationships; however, it can be highly challenging to conduct biological experiments on a large number of genes due to limited lab resources. On the other hand, computational resources are relatively cheaper; however, reliably identifying causal relationships is known to be an NP-hard problem.^67^ The described iterative algorithm takes advantage of both sides by improving the set of computationally identified genes utilizing experimentally learned causal relations.

We conjecture that DECODER will become even more powerful in the near future as the number of AD studies providing brain gene expression and neuropathology data continues to increase. Many of these studies also provide multi-omic data, which will let us utilize a higher sample size for learning regulatory multi-omic module networks for identified biomarker genes.

## Methods

### Gene expression and neuropathology datasets

We used data from the ROSMAP,^12,13^ ACT,^15,21^ and MSBB studies. Around half of the people in each cohort had been diagnosed with dementia by the time of death. MSBB neuropathology data was made available by the AMP-AD Knowledge Portal of Sage Bionetworks through https://www.synapse.org/ with Synapse ID syn6101474. We accessed neuropathology data from the ROSMAP and ACT studies through data use agreements. ACT neuropathology data includes four types of Aβ and tau quantifications: (1) immunohistochemistry (IHC) measured on fresh-frozen brain tissue, (2) IHC measured on formalin-fixed, paraffin-embedded (FFPE) brain tissue, (3) histelide^66^ quantification from FFPE slides, and (4) Luminex. We chose histelide for Aβ and IHC FFPE for tau since these two led to the strongest average association with gene expression levels.

ROSMAP RNA-Seq data and MSBB RNA-Seq data were made available by Sage Bionetworks on the AMP-AD Knowledge Portal^11^ with Synapse IDs syn3505732 and syn7391833, respectively. The ACT RNA-Seq data^15^ was collected by the Allen Institute for Brain Science, Kaiser Permanente Washington Health Research Institute (KPWHRI), and the University of Washington (UW), and it was made available on http://aging.brain-map.org. We used normalized and log-transformed RNA-Seq read counts for all datasets.

ACT RNA-Seq data is part of the larger ACT project — a collaboration between KPWHRI and UW — that collected clinical and genetic data from thousands of individuals. The approximately 100 ACT individuals from whom we have RNA-Seq measurements were specifically selected for a traumatic brain injury (TBI) study; half of the selected individuals sustained a TBI with loss of consciousness during their lifetime. As a result, the RNA-Seq cohort does not necessarily reflect the demographics of the entire set of thousands of ACT individuals. Therefore, the ACT study assigned each RNA-Seq sample a weight based on the demographic information of each sample in the TBI study. Details of this weighting scheme and the weights themselves are provided on http://aging.brain-map.org. As recommended by the ACT study, we took the weights into account when computing gene expression-neuropathology associations for the ACT data so that the samples reflect the original ACT cohort demographics.

### Probabilistic modeling of the relation between neuropathology and gene expression levels

Let us assume that for each of *R* brain regions, we observe a data matrix 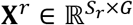 that contains the gene expression measurements on *G* genes and for *S_r_* samples (i.e., individuals), and we also observe a neuropathology feature for the same *S_r_* samples, represented by a column vector 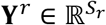. Let us also assume that **X**^*r*^ and **Y**^*r*^ are standardized so that each column in each matrix has a mean of 0 and variance of 1.

We model the data observed for the brain region *r* using the following generative model, where 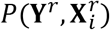 represents the joint probability distribution of the neuropathology feature and the expression level of gene *i*, both measured in that region:

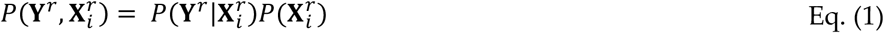

Assuming a Gaussian distribution for the conditional 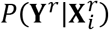 and a uniform prior distribution over 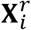:

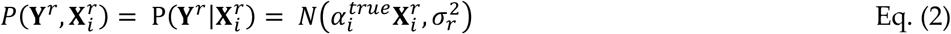

where *i* is a gene index and *r* is a region index, and 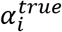 is a parameter that represents the true association we want to learn between the expression level of gene *i* and the neuropathology feature.

Using the probability density formulation of Gaussian distribution and assuming independent samples:

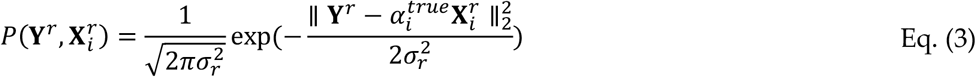

where 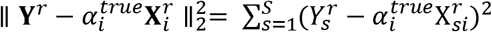 represents the squared Euclidean norm of 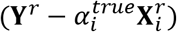. Assuming the brain regions are also independent of each other:

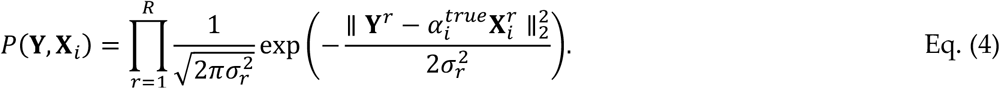

Therefore:

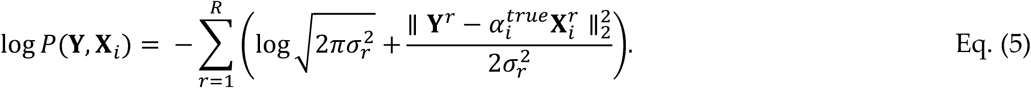

To learn the true association 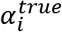 between the expression level of gene *i* and the neuropathology feature based on this probabilistic model, we maximize the joint log-likelihood, which corresponds to the following optimization problem:

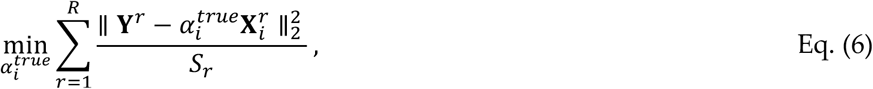

assuming that 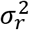 is proportional to *S_r_* so that the standard error of the mean is the same for each region independent from the number of samples measured for that region. This is a reasonable assumption because a highly sampled brain region may not be the one that is highly relevant to AD. When we take the derivative of this objective function with respect to 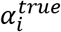 and set it to zero to get the 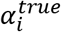 minimizing it, we get:

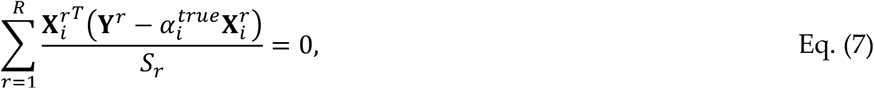

which gives below solution for 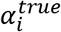:

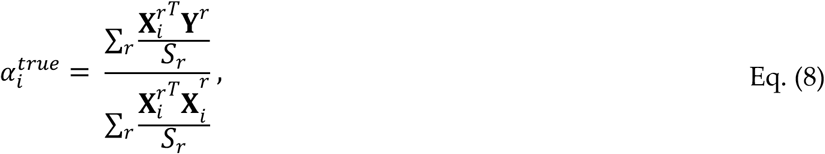

and so:

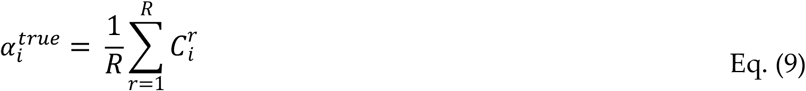

where 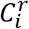 represents the association of the expression of gene *i* with the neuropathology in region *r* (i.e., the estimate of the Pearson’s correlation coefficient (PCC) between 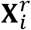 and **Y**^*r*^). Eq. (9) holds because 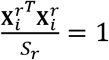 and 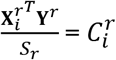 since both 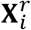 and **Y**^*r*^ are standardized. Thus, the true association between the expression level of each gene *i* and the neuropathology feature can be computed as the equally weighted average of the individual region-based associations for that gene. The resulting **α**^*true*^ values correspond to the scores obtained by our equal weights score (EWS) approach.

As Eq. (9) shows, EWS computes the true association for each gene *i* as the mean of the regional associations for that gene. Thus, it equally weighs each region while computing the true gene-neuropathology associations, where each weight value is equal to 1/*R*. Figure 12A represents EWS, where region weights are assumed to be equal; hence, the final gene scores are computed as the average across input region associations.

**Figure 12.**
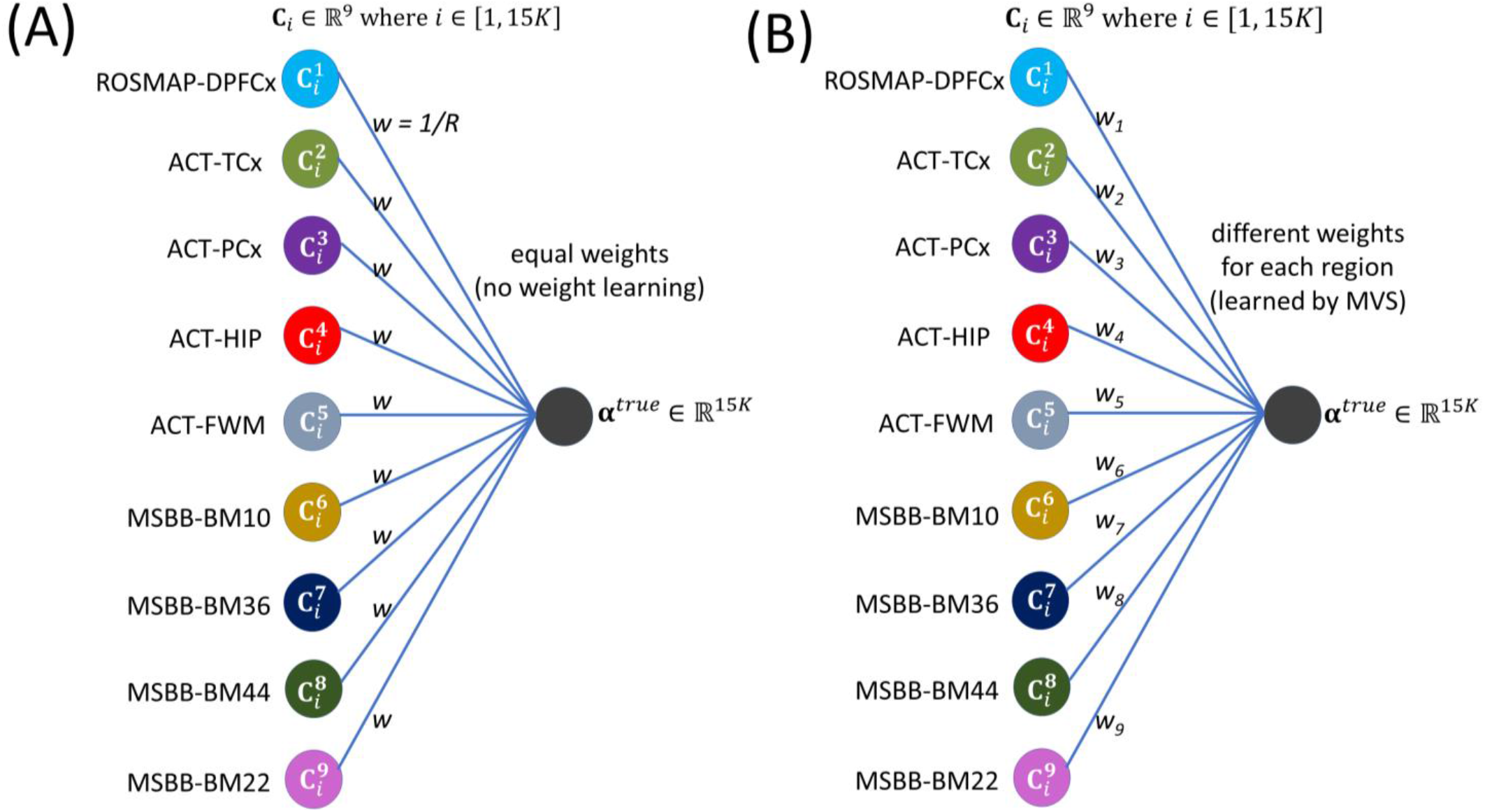
EWS and MVS representations. The single hidden unit represents **(A)** EWS and **(B)** MVS. Input node region colors are matched to region colors in Figure 1. An edge between two units represents the weight of a feature in computing the linear combination of feature values to compute the final concordance-based scores.

However, different regions are likely to have a different importance in AD; rather than assuming equal weights, we may want to learn the weights *w_r_* for each region *r* while computing the true underlying association 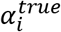 for each gene. Therefore, our maximum variance score (MVS) approach assumes that the true neuropathology association for each gene *i* is a weighted mean of the individual region-based associations for that gene:

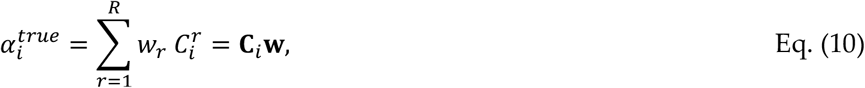

where **C**_*i*_ represents a row of 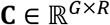, a matrix representing the individual region-based PCCs between genes and neuropathology, and 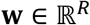 is a unit column vector representing region weights (Figure 12B).

There are multiple ways to assign values to *w_r_*. One natural option is to choose *w_r_* values so the learned **α**^*true*^ represents as much information about the regional neuropathology associations as possible (i.e., we maximize the variance). That is how MVS learns the weight values.

Without loss of generalization, we assume that the true association vector **α**^*true*^ has a mean of 0 across all genes 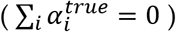. Thus, maximizing the variance of the data through **α**^*true*^ corresponds to the following optimization problem:

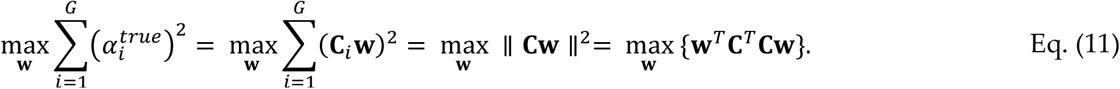

Since **w** is a unit vector, **w**^*T*^**w** = 1, the preceding optimization problem is equivalent to

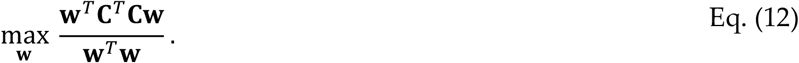

Since **C**^*T*^**C** is a positive semi-definite matrix, the **w** vector that maximizes the quantity 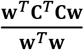 (also known as the Rayleigh quotient) is the eigenvector corresponding to the maximum eigenvalue of **C**^*T*^**C**. Thus, **α**^*true*^ = ***C*w** corresponds to the first principal component resulting from applying principal component analysis (PCA) applied to the individual region associations matrix **C**. The resulting **α**^*true*^ values correspond to the scores obtained by our MVS method.

The MVS objective while learning **w** (i.e., the weights of the individual regions in Eq. (10)) is to capture the maximum variance in the input associations by projecting them onto the direction of **α**^*true*^. Another objective would be to reconstruct input associations from **α**^*true*^ (by multiplying **α**^*true*^ back by **w**^*T*^) as accurately as possible (i.e., with a minimal error between the input associations and the reconstructed associations). Then the objective becomes:

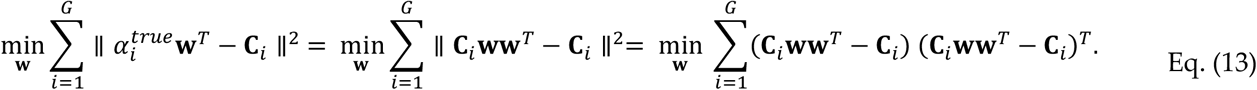

This objective has been shown to lead to the same **w** vector as that learned by MVS.^68^

Each of EWS and MVS learns a *linear* embedding of the input data (i.e. PCCs from the individual regions) to a single-dimensional vector. Non-linear embedding might be useful since it lets us capture more complex relationships between input variables (i.e., brain regions) and the learned low-dimensional space. Therefore, we developed our best reconstruction score (BRS) approach, which aims to reconstruct the input with a minimal error using an artificial neural network architecture. Compared to Eq. (13) above, BRS uses an additional set of weights (i.e., an additional hidden layer in the network) and applies a nonlinear activation function on data supplied to each hidden layer *h* (Figure 13). The final BRS objective is:

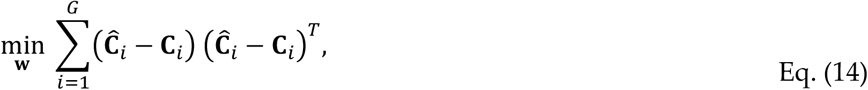

**Figure 13.**
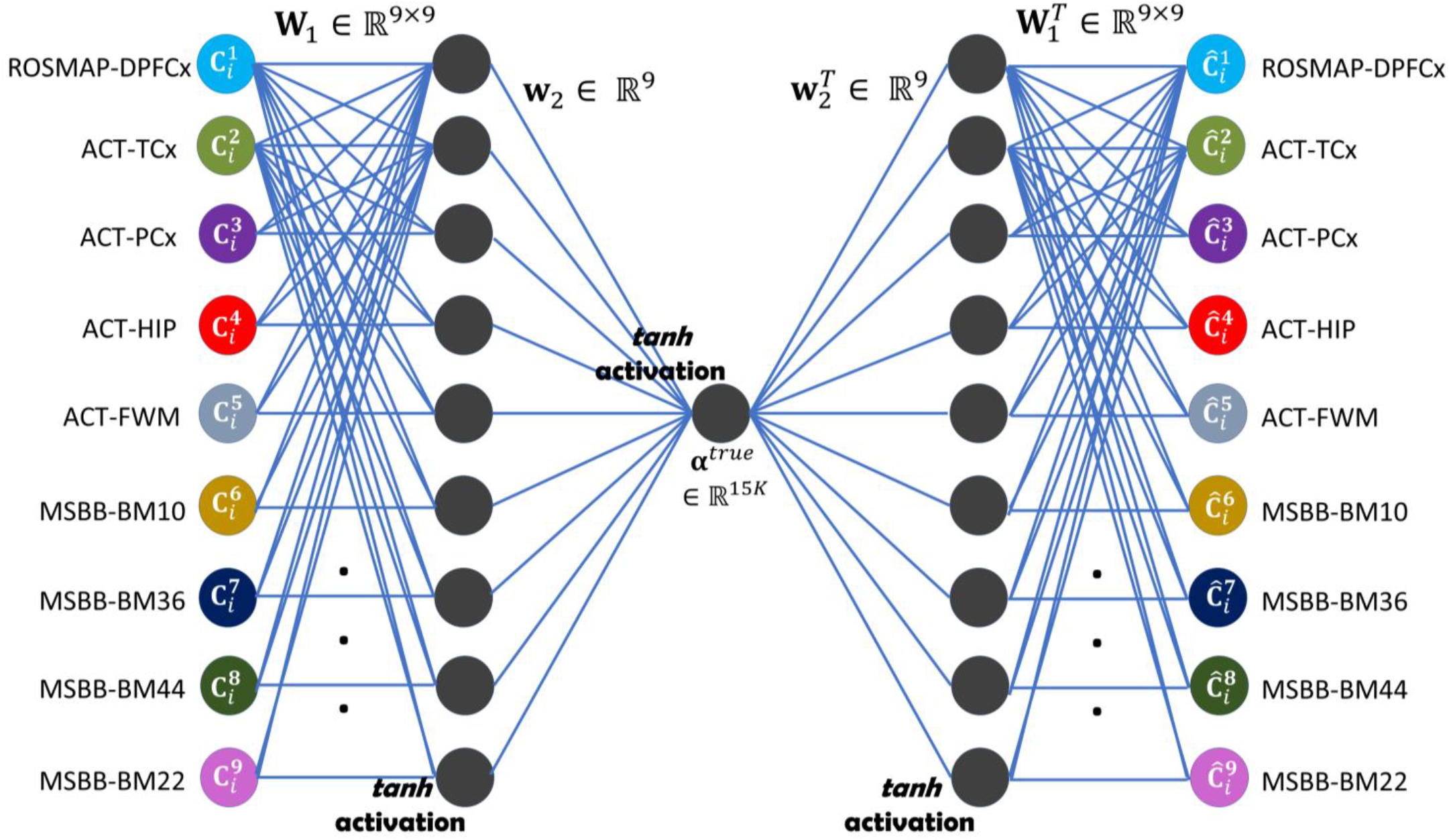
The BRS representation. We used a neural network architecture with two hidden layers and we used the nine individual region associations as input. Input node region colors are matched to region colors in Figure 1. Hidden units are shown in gray. An edge between two units represents the weight of a feature in computing the linear combination of the feature values to be passed to the activation function. The single hidden unit in the final hidden layer (bottleneck layer) represents the BRS values of the genes.

where 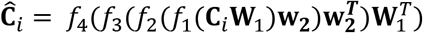 represents the data reconstructed by BRS (i.e., the reconstructed PCCs). As demonstrated in Figure 13, we use hyperbolic tangent function *fanh*(.) as *f*_1_, *f*_2_, and *f*_3_, and we use *R* (=9) units in the intermediate hidden layer. Thus, 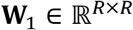 is a 9 × 9 matrix, and 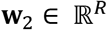 is a column vector of nine elements like **w** in MVS. We use an identity activation function in the output layer (i.e., *f*_4_(*x*) = *x*) and the mean-squared error (MSE) to measure reconstruction quality since gene expression-neuropathology associations (**C**) are continuous-valued between -1 and 1. Our resulting BRS network resembles an autoencoder network architecture, and the three dots in each of the encoder and decoder networks in Figure 13 imply full-connectivity in the intermediate layer.

Unlike EWS or MVS, the cost function for BRS is not convex. A common approach to learning **W**_1_ and **w**_2_ is to start with some random weight values and iteratively forward/backward propagate through the network (Figure 13) to update weight values.^69^ In each iteration, the reconstruction error is computed first by forward propagation; gradients that reduce the error are then computed through back-propagation; and weights are updated according to those gradients. Since the cost function of BRS is non-convex, the resulting weights may depend on the weight values at initialization. Therefore, we performed 100 BRS runs initialized with different random weights and averaged the results across these runs.

### Data collection and association analysis for the experiments with TCGA data

To examine cancer survival associations, we used clinical data and RNA-Seq data provided by the Broad Institute (http://gdac.broadinstitute.org/). We used the “SurvCorr”^70^ R package available on CRAN to estimate a correlation between a right-censored survival time and a gene expression level. Table 7 lists the expansions for cancer type abbreviations we used in our experiments with TCGA data:

**Table 7.** Cancer type abbreviations.

|  |  |
| --- | --- |
| ACC | Adrenocortical carcinoma |
| BLCA | Bladder urothelial carcinoma |
| BRCA | Breast invasive carcinoma |
| CESC | Cervical and endocervical cancers |
| CHOL | Cholangiocarcinoma |
| COAD | Colon adenocarcinoma |
| DLBC | Lymphoid neoplasm diffuse large B-cell lymphoma |
| ESCA | Esophageal carcinoma |
| GBM | Glioblastoma multiforme |
| HNSC | Head and neck squamous cell carcinoma |
| KICH | Kidney chromophobe |
| KIRC | Kidney renal clear cell carcinoma |
| KIRP | Kidney renal papillary cell carcinoma |
| LAML | Acute myeloid leukemia |
| LGG | Brain lower grade glioma |
| LIHC | Liver hepatocellular carcinoma |
| LUAD | Lung adenocarcinoma |
| LUSC | Lung squamous cell carcinoma |
| MESO | Mesothelioma |
| OV | Ovarian serous cystadenocarcinoma |
| PAAD | Pancreatic adenocarcinoma |
| PCPG | Pheochromocytoma and paraganglioma |
| PRAD | Prostate adenocarcinoma |
| READ | Rectum adenocarcinoma |
| SARC | Sarcoma |
| SKCM | Skin cutaneous melanoma |
| STAD | Stomach adenocarcinoma |
| TGCT | Testicular germ cell tumors |
| THCA | Thyroid carcinoma |
| THYM | Thymoma |
| UCEC | Uterine corpus endometrial carcinoma |
| UCS | Uterine carcinosarcoma |
| UVM | Uveal melanoma |

### Identification of C. elegans orthologs

To enable biological testing of the human genes identified using our computational analysis, we obtained the RBHs between human and *C. elegans*. To do so, we first identified all unique protein sequences for each potential marker gene using the “biomaRt”^71^ R package available on CRAN. Then, we used the latest version of the NCBI BLAST tool^72^, which we downloaded from ftp://ftp.ncbi.nlm.nih.gov/blast/executables/blast+/LATEST, version 2.6.0 as of 8/15/17, to identify the *C. elegans* orthologs for each complete human protein query sequence. We downloaded the *C. elegans* protein sequences from http://www.wormbase.org/species/c_elegans. We took into account only the protein pairs mapped from human to *C. elegans* with a BLAST *e*-value smaller than 10^-30^. For each *C. elegans* isoform, we identified the corresponding human genes, again using the NCBI BLAST tool, and used only the orthologs that achieved a BLAST *e*-value smaller than 10^-30^. This process resulted in high-confidence RBHs for us to test in *C. elegans*.

### C. elegans cultivation and RNAi treatment

Experimental worm populations of GMC101 animals were obtained from the Caenorhabditis Genetics Center (CGC) and cultivated on NGM plates with OP50 at 15C.^33,73^ Care was taken to ensure that the animals were never starved and the plates remained free of contamination.

The gene-specific RNAi clones were obtained from the commercial Ahringer or Vidal *C. elegans* RNAi-feeding libraries (BioScience, Nottingham, UK). Each bacterial clone was struck-out onto LB plates containing carbenicillin (50 ug/ml) and tetracycline (10 ug/ml). Single colonies were then seeded into 5 ml LB + carbenicillin (50 ug/ml) and tetracycline (10 ug/ml) for growth overnight on a 37C rotator. 100 ul of each overnight culture was then inoculated into 10 ml of LB containing carbenicillin (50 ug/ml) and tetracycline (10 ug/ml) and IPTG (5mM) and incubated on a 37C rotator for 4 hours. Each bacterial growth was then centrifuged at 3500 X G for 25 minutes, decanted, and the pellet resuspended in 0.5 ml of LB containing carbenicillin (50 ug/ml), tetracycline (10 ug/ml), and IPTG (5 mM). To verify RNAi conditions’ plasmid DNA, each RNAi clone was purified and assessed through PCR (polymerase chain reaction) with sequence-specific primers or through Sanger sequencing.

### Nematode paralysis assays

Paralysis assays were performed by visually inspecting animals daily to determine if they were capable of spontaneous movement or if they were paralyzed. A robotic system equipped with a digital camera was used to obtain images of individual wells of a 12 well-plate at 5-minute intervals over the entire course of the experiment. Each well contained 30-40 individual animals expressing Aβ. Through analysis of serial images from each plate, the age at which each animal stopped moving could be easily determined. We applied this system to the transgenic Aβ model line GMC101 to determine the time of paralysis onset for each individual animal. Paralysis data was plotted using Oasis2. Statistical significance of mean paralysis time-points between RNAi conditions was determined by a weighted log-rank^74,75^ test.

Prior to loading on the experimental plates, animal populations were amplified on high-growth plates seeded with NA22 bacteria. Worm populations were developmentally synchronized by hypochlorite treatment, and the remaining eggs were deposited on unseeded plates overnight. Synchronized larval stage 1 animals were washed off unseeded plates and moved onto standard *C. elegans* RNAi plates containing carbenicillin (50 mg/ml), tetracycline (10 mg/ml), and IPTG (5 mM) 48h at 20C. These developmentally synchronized, late larval stage 4-populations were then washed and transferred to their respective RNAi conditions on 12-well plates. We used standard RNAi conditions plus FuDR (100 ug/ml) to prevent progeny and neomycin (200 mg/ml) to prevent fungal growth.^76^ Each RNAi condition was tested in 2-3 wells as technical replicates. At least three biological replicates, each started on different weeks, were conducted for each RNAi clone.

### Oxygen consumption assay

We quantified the oxygen consumption rates of day-two adult worms cultivated from L1 on Y53G8AL.2 RNAi or the EV control RNAi at 20C. Oxygen consumption was measured utilizing the Seahorse X24 Bioanalyzer (Seahorse Biosciences, MA, USA).^54^ Briefly, young adult worms were washed from NGM plates and rinsed from RNAi bacteria with M9 buffer. Approximately 40-50 worms were pipetted into each well in Seahorse XF24 Cell Culture Microplates, with the final volume of 500μL M9. The number of worms in each well was quantified. Basal respiration was analyzed using the average respiration of five technical replicates per condition, and eight recordings over the course of an hour were made of each condition. Three independent biological replicates were conducted on different weeks. For each experiment, we compared the relative rate of oxygen consumption between the paired conditions. In each plate, nematode-free wells were used as control. Oxygen consumption rates were then normalized by the number of worms. All three replicates showed that the animals cultivated on the Y53G8AL.2 RNAi consumed about half as much oxygen as those grown on the control RNAi bacteria (Figure 9).

### Multi-omic module network learning

Given a matrix of values for each variable-sample pair from a number of variables and a number of samples, MGL iteratively learns an assignment of the variables to a user-supplied number of modules and the conditional dependence network among modules. In a conditional dependence module network, two modules that are disconnected are independent of each other given all other modules.

For the MGL algorithm to run in a reasonable amount of time, from the 420,132 DNA methylation sites in the ROSMAP data, we selected 20,132 DNA methylation sites with the highest variance across samples. Supplementary Figure 3 shows the standard deviation of all methylation sites; the red vertical line represents the standard deviation threshold we used to select the methylation sites to use in MGL. In addition to DNA methylation levels, we used expression levels of 14,912 protein-coding genes and 309 miRNAs from the ROSMAP study. Thus, we learned a network of modules from a total of 35,354 variables. We set the module count to 500 before running the algorithm. This resulted in an average of 70 genes per module, while the variable count varied across modules. Since MGL has a non-convex objective function, we performed five runs of the MGL algorithm with different initial parameter values and selected the run with final parameter values leading to the highest likelihood of the training data. Supplementary Table 3 shows information about each of the 500 modules learned by MGL. We note that since the dependence network MGL algorithm^40^ learns is based on an inverse covariance matrix, a positive edge weight shown in column (J) of Supplementary Table 3 corresponds to an up-regulation, and a negative edge weight corresponds to a down-regulation.

## Acknowledgments

This work was supported by National Science Foundation (NSF) grants DBI-1355899 and CAREER DBI-1552309 to SL; National Institutes of Health (NIH) grant AG049196-01A1 to SL and SC; NIH grants P30AG013280 and P50AG005136 to MK; and NIH grant F32AG054098-01S1 to JR. We are grateful to Mount Sinai/JJ Peters VA Medical Center Brain Bank for making the MSBB gene expression and neuropathology data available through the AMP-AD Knowledge Portal; to Caenorhabditis Genetics Center (CGC) for the worm reagents; and to Nick Terzopoulos for preparing the nematode cultivation plates. Generation of the ACT RNA-Seq data was funded by a grant to CD Keene, RG Ellenbogen and Ed Lein from the Paul G. Allen Family Foundation, and supported by NIH grants U01AG006781 and P50AG005136 and the Nancy and Buster Alvord Endowment. ROSMAP data collection was supported through funding by the National Institute on Aging (NIA) grants P30AG10161, R01AG15819, R01AG17917, R01AG30146, R01AG36836, U01AG32984, U01AG46152; the Illinois Department of Public Health; and the Translational Genomics Research Institute (TGen).

